# A biocatalytic platform for the production of substituted 2-quinolones and (thio)coumarins

**DOI:** 10.1101/2024.12.20.629698

**Authors:** Nika Sokolova, Angelina Osipyan, Lili Zhang, Matthew R. Groves, Sandy Schmidt, Kristina Haslinger

**Affiliations:** Department of Chemical and Pharmaceutical Biology, University of Groningen, The Netherlands

## Abstract

2-quinolones are privileged scaffolds for drug discovery that are relatively rare in nature. Here, we characterise two promiscuous fungal polyketide synthases AthePKS and FerePKS, which we had previously found to produce quinolones *in vitro*. We challenged the enzymes with several substituted anthranilic acid derivatives, revealing their ability to produce precursors of pharmaceutically relevant quinolones. We also discovered that AthePKS and FerePKS accept other 2-substituted benzoic acids, leading to the formation of coumarin and thiocoumarin scaffolds. We applied AthePKS in an artificial enzymatic cascade towards an antimicrobial 4-methoxy-1-methyl-2-quinolone and demonstrated its *in vivo* feasibility by successfully expressing the pathway in *Escherichia coli*. Lastly, we determined the crystal structure of AthePKS, suggesting hotspots for enhancing its catalytic efficiency by enzyme engineering. Our results provide a framework for further engineering of enzymatic routes towards privileged heteroaromatic scaffolds and derivatives thereof.

## Introduction

Natural products (NPs) are an inexhaustible source of biologically active compounds. Structurally and functionally refined through evolution, these molecules have been a major source of inspiration in drug discovery and development for decades^1,2^. However, the direct use of NPs is often impeded by their poor pharmacokinetics, while structural complexity renders chemical modification challenging^3,4^. To bypass these limitations, many researchers have focused on utilising the basic molecular scaffolds of NPs as platforms for drug discovery^5^.

One such “privileged” scaffold is 2-quinolone, also known as carbostyril or 1-azacoumarin. Found in a group of bioactive alkaloids mainly from Rutaceae plants, it is also at the core of several marketed drugs with different modes of action (Figure 1). Many synthetic 2-quinolone derivatives have also been shown to possess promising antimicrobial^6–8^, anti-Alzeimer’s^9^, anticancer^10^ and anti-diabetic^11^ activities. Due to the intrinsic photoactive properties of the conjugated 2-quinolone system, this scaffold is also investigated for use in luminescent materials and cationic metal sensors, among other functional organic materials^12^.

**Figure 1.**
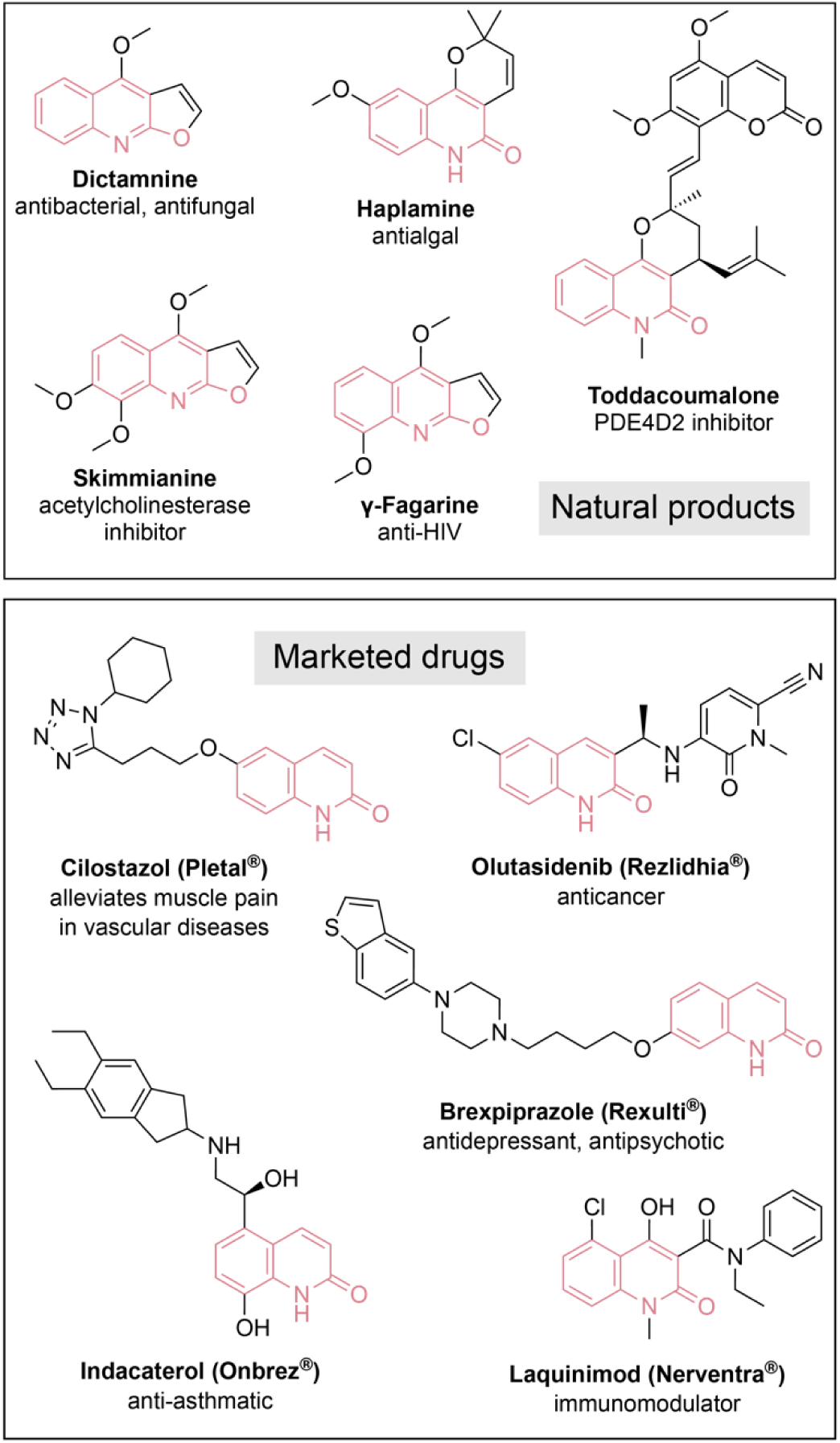
Some of the bioactive natural products and marketed drugs containing the 2-quinolone scaffold.

Numerous strategies for the chemical synthesis of 2-quinolones have been developed^12,13^. In nature, this scaffold is typically represented by 4-hydroxy-1-methyl-2-quinolone, as synthesised by type III polyketide synthases (T3PKSs). The reaction involves decarboxylative condensation of malonyl-coenzyme A (CoA) with *N*-methylanthraniloyl-CoA to form a diketide intermediate which then cyclises to yield the heteroaromatic quinolone. However, only a handful of quinolone synthases, either from plants or bacteria, have been characterised to date^14–17^, and their application in the biosynthesis of quinolone derivatives remains underexplored.

In a previous study^18^, we performed activity profiling of 37 fungal T3PKS with 12 acyl-coenzyme A substrates. Unexpectedly, half of the active enzymes converted *N*-methylanthraniloyl-CoA to the heteroaromatic quinolone. From this set of active enzymes, we selected two best-performing T3PKSs for further characterisation in this study. We used the enzymes in combination with a bacterial CoA ligase to generate a series of substituted quinolone derivatives. We also demonstrate the ability of both enzymes to accept (thio)salicylic acid derivatives, leading to the formation of three distinct heteroaromatic scaffolds. We harnessed this promiscuity to develop a biocatalytic cascade towards an antimicrobial O-methylated quinolone for the first time and demonstrated its feasibility in a microbial host. The functional and structural insight generated in this study will guide future efforts towards sustainable production of privileged heteroaromatic scaffolds.

## Results and discussion

### AthePKS and FerePKS produce N-, O- and S-containing heterocycles *in vitro*

First, we set out to test the substrate scope of the two T3PKSs on several anthranilic acid derivatives. Since T3PKSs typically act on CoA-activated carboxylic acids, we employed the anthranilate-CoA ligase PqsA from *Pseudomonas aeruginosa*^19^ (UniProt: Q9I4X3) to generate the substrates for the T3PKSs. To streamline the analysis, we opted to perform the assays in a one-pot continuous cascade with the respective carboxylic acids (**1**-**8**) as starter substrates and malonyl-CoA (**9**) as the extender unit. We first tested the viability of such an approach with *N*- methylanthranilic acid (**1**) using 1 mol% of each enzyme. Under these conditions, PqsA-AthePKS and PqsA-FerePKS converted 12-24% of **1** through *N*-methylanthraniloyl-CoA (**10**) to the corresponding quinolone (**12**, Supplementary Figure 1a and 2a).

We then turned to substituted anthranilic acid derivatives **2**-**5** and assessed first whether they could be accepted by PqsA (Supplementary Figure 3). We observed full conversion of **2** and **3** to the CoA-activated intermediate, whereas **4** and **5** were only partially converted (12 and 36%, respectively). In the continuous cascade, the non-*N*-methylated anthranilic acid (**2**) was then converted to the quinolone product by both AthePKS and FerePKS, albeit less efficiently than **1** (Figure 2a, Supplementary Figure 2). 5-Chloro-anthranilic acid (**3**) was accepted better than **2** by both enzymes. Despite the low activity of PqsA with substrate **4**, we detected trace amounts of the putative product with the expected *m/z* of 178 in reactions with AthePKS. 3,4- Dimethoxyanthranilic acid (**5**) was not converted by either enzyme.

**Figure 2.**
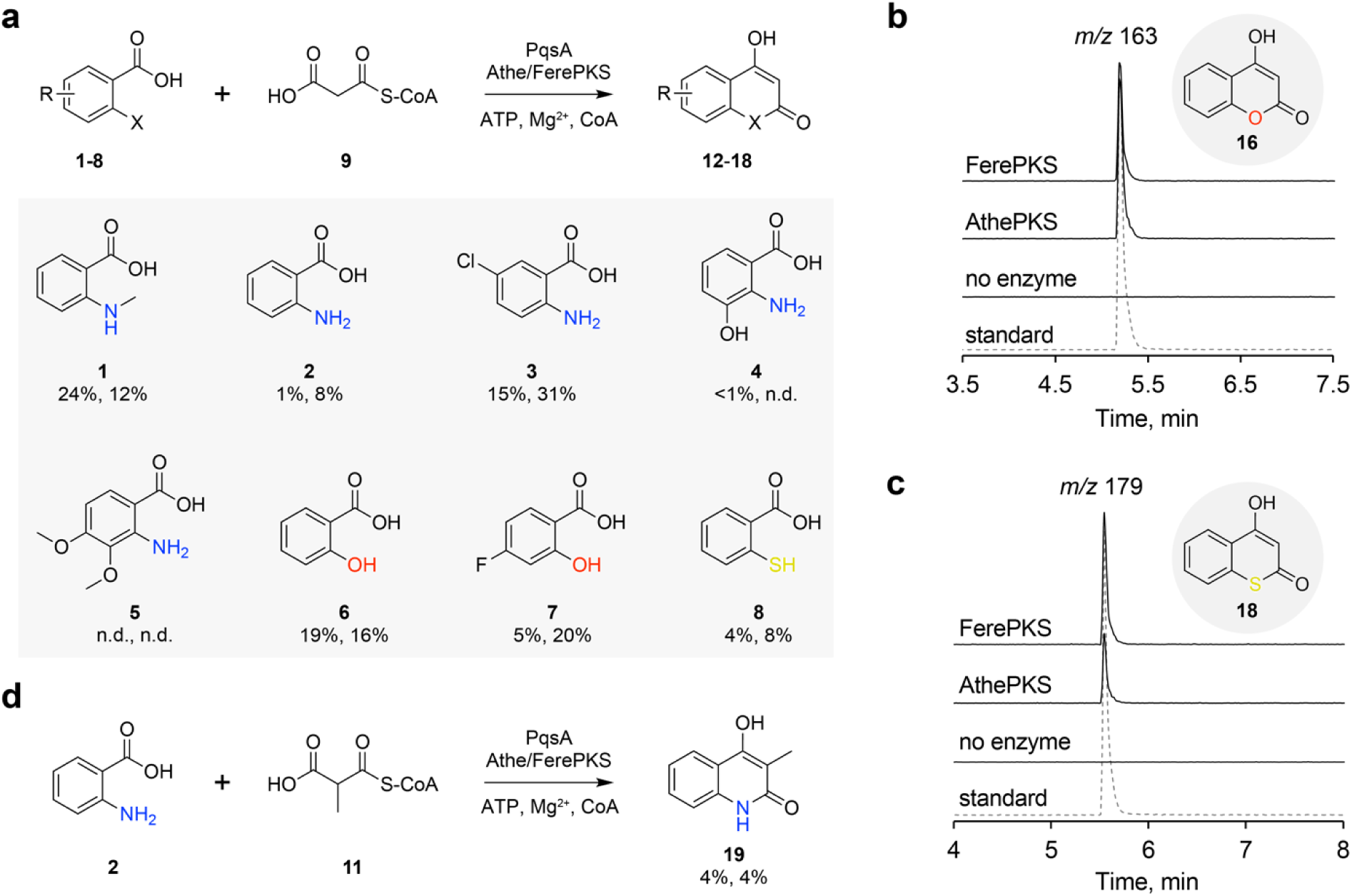
Substrate scope analysis of AthePKS and FerePKS. a) One-pot continuous cascade with PqsA, Athe/FerePKS, extender unit **9** and carboxylic acids **1-8** yields products **12-18**. The percentages correspond to % conversion of a given substrate under the assay conditions by AthePKS and FerePKS, respectively; “n.d.” - not detected. b) Formation of 3-methyl-4-hydroxy-2-quinolone (**19**) from **2** and extender unit **11** by PqsA and Athe/FerePKS. c-d) Analysis of the reaction products of PqsA and Athe/FerePKS with substrates **6** and **8** showing the formation of coumarin and thiocoumarin scaffolds by Liquid Chromatography-coupled mass spectrometry (LC-MS). Dotted lines represent the authentic standards of 4-hydroxycoumarin (**16**) and 4-hydroxythiocoumarin (**18**).

We then challenged the enzymes with other 2-substituted benzoic acids that we hypothesised would lead to heterocycle formation. Although CoA activation by PqsA was somewhat lower for substrates **6**-**8** (74, 45, and 79% respectively, Supplementary Figure 3), they were accepted both by AthePKS and FerePKS with conversions comparable to those with anthranilic acid derivatives **1-3**, leading to the formation of several 4-hydroxy(thio)coumarin derivatives (Figure 2b-c). An overview of products **12**-**18** is presented in Supplementary Figure 1b. Lastly, we assessed whether AthePKS and FerePKS could incorporate methylmalonyl-CoA (**11**) as an extender unit, as observed in nature for several fungal T3PKS. We incubated both enzymes with substrate **2** and **11** to yield **19** – a C-methylated quinolone with reported antimicrobial activity against several species of *Mycobacterium*^20^ (Figure 2d).

To our knowledge, this is the first report of the enzymatic formation of quinolones **14**-**15**, coumarin **17** and thiocoumarin **18**. The 6-chlorinated quinolone scaffold of **14** resembles the common pharmacophore of HIV-1 non-nucleoside reverse transcriptase inhibitors^21^, which is also found in the anticancer drug Olutasidenib (Figure 1). The 8-hydroxylation of **15** is a common motif in several clinical trial drugs such as Abediterol and Navafenterol; it is also common in furoquinoline alkaloids from plants, where it is often methylated. 4-Hydroxycoumarin **16** is the direct precursor of Warfarin and related anticoagulant compounds, while its fluorinated analogue **17** is the monomer of a potent homodimeric inhibitor of human NAD(P)H quinone oxidoreductase-1 (NQO1), an enzyme overexpressed in several types of tumour cells^22^. Thiocoumarins, including **18**, possess anticoagulant properties similar to coumarins and have been used in commercial rodenticides^23^.

Thus, even without optimisation, the one-pot cascade with PqsA and a fungal T3PKS yielded a variety of heteroaromatic pharmaceutical intermediates with moderate to good conversions. We then sought to further extend the cascade towards a more complex bioactive product in order to demonstrate its applicability.

### Screening of methyltransferases for the 4-OH-methylation of quinolones

While researching the diversity of natural 2-quinolones from *Rutaceae* plants, we encountered 4- methoxy-1-methyl-2-quinolone (**20**), a compound with reported antimicrobial^24,25^, antimalarial^26^ and anti-HIV^27^ activity. Compound **20** also appears to be the precursor of more complex *Rutaceae* alkaloids, although their biosynthesis is not experimentally elucidated (Figure 3a). It has not escaped our notice that **20** is just one enzymatic step away from the AthePKS product **12** – specifically, it is methylated at the 4-hydroxy position (Figure 3b). We therefore hypothesised that AthePKS could be applied in a biocatalytic cascade to produce **20**.

**Figure 3.**
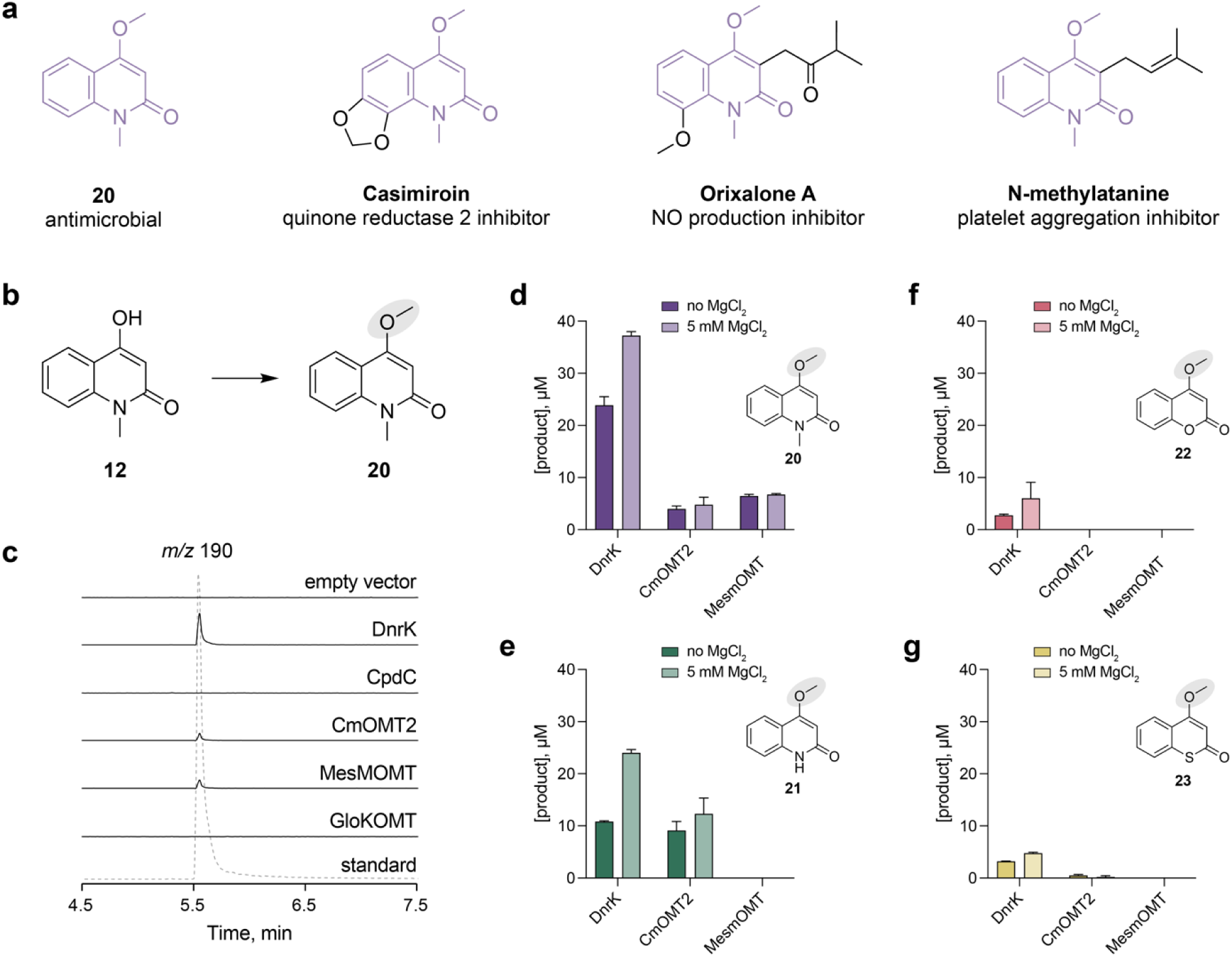
Screening of O-methyltransferases (OMTs) for the production of **20**-**23**. a) Chemical structures of 4-methoxy-1-methyl-2-quinolone (**20**) and quinolones hypothesised to be derived from it. b) 4-OH methylation of **12** to yield **20**. c) Screening of five OMTs for the methylation of **12** in cell-free extracts. Solid lines represent the extracted ion chromatograms in low-resolution LC-MS of the expected reaction product; dotted line represents the chemically synthesised reference of **20**. d-g) Assays with purified DnrK, CmOMT2 and MesMOMT for the methylation of quinolones **12** and **13**, coumarin **16** and thiocoumarin **18**. The reactions were incubated with 0.5 mM substrate, 2 mM SAM and 5 µM enzyme for 24 h.

Since the biosynthesis of **20** is not elucidated and, to our knowledge, no heterologous *O*- methyltransferase (OMT) was ever reported to catalyse the methylation of **12**, there was no obvious candidate enzyme for this biosynthetic step. Hence, we searched the literature for OMTs that are active on substrates that are structurally similar to **12** and identified three candidates: the bacterial enzyme DnrK from the biosynthetic pathway of daunorubicin^28^, the plant enzyme CmOMT2 that methylates hydroxycoumarins in other positions^29^, and the fungal enzyme CpdC from the biosynthetic pathway of calipyridone A^30^ (Supplementary Figure 4). We also noticed that the enzymes MesMOMT and GloKOMT from our previous study on the methylation of catechols^31^ share the same structural fold as the three aforementioned OMTs, and included them in the screening (Supplementary Figure 5).

We expressed the five genes in *E. coli* BL21(DE3) and tested the activity of the OMTs with **12** and *S*-adenosylmethionine (SAM) as methyl donor in cell-free lysates. With three out of five enzymes, we detected a product with an *m/z* value that corresponds to mono-methylated **12** and co-elutes with a chemically synthesised reference of **20** (Figure 3c). Under the expression conditions, DnrK yielded the most **20**. Interestingly, MesMOMT, which methylated caffeic acid and other catechols in our previous study^31^, was also active on **12**.

We expressed and purified the three active OMTs and tested them with **12** at 1 mol% enzyme to substrate. DnrK was still the best-performing enzyme, yielding 24 µM **20**, which led us to choose this enzyme for the cascade applications. Interestingly, the addition of 5 mM MgCl_2_ to the reaction mixture increased the product concentration by 35% (Figure 3d). DnrK was also active with other heterocycles **13**, **16** and **18**, albeit the conversions were lower than that of **12** (Figure 3 e-g, Supplementary Figure 6).

### Engineering a biocatalytic cascade towards an antimicrobial 4-methoxy-1- methyl-2-quinolone

With the necessary enzymes at hand, we proceeded to assemble a biocatalytic cascade towards **20** from **1** (Figure 4a). The initial conditions for the cascade were adapted from the PqsA reaction^19^ with the addition of AthePKS and DnrK at the same molar concentration as PqsA, and the corresponding cofactors. The three-step one-pot continuous cascade yielded 17 ± 0.75 µM **20** from 500 µM substrate after 24 h (Figure 4b). We then sought to optimize several parameters to increase the titre of the final product. We found that buffering the reaction mix with Tris instead of HEPES at a lower pH is beneficial for the production of **20** (Figure 4c). We also found that increasing the concentration of Mg^2+^ above 5 mM in the reaction mix does not improve the titre of **20** but drives the CoA ligation to completion (Figure 4d). Next, we optimised the starting concentration of CoA in the cascade. During the T3PKS reaction, CoA is released from **9** and **10** as a byproduct and can in principle be recycled by the CoA ligase. Therefore, we tested the lower limits of the starting CoA concentration and found that adding as little as 15.6 µM CoA does not drastically affect the final titre of **20**, and a 1:8 CoA-to-substrate ratio afforded the highest conversion to **20** (Supplementary Figure 7). Lastly, we evaluated the effect of altering the molar ratio of the enzymes in the cascade. As expected, increasing the concentration of AthePKS and DnrK relative to PqsA improved the product titre (Figure 4e). After optimisation, the concentration of **20** produced by the cascade increased from 17 ± 0.75 µM to 177 ± 2 µM (35% conversion of **1**). The conversion further increased to 41% upon incubation for 72 h (Figure 4f). When the substrate load was increased to 1000 µM, the reaction was completed after 48 h and the highest conversion was 25% (Supplementary Figure 8).

**Figure 4.**
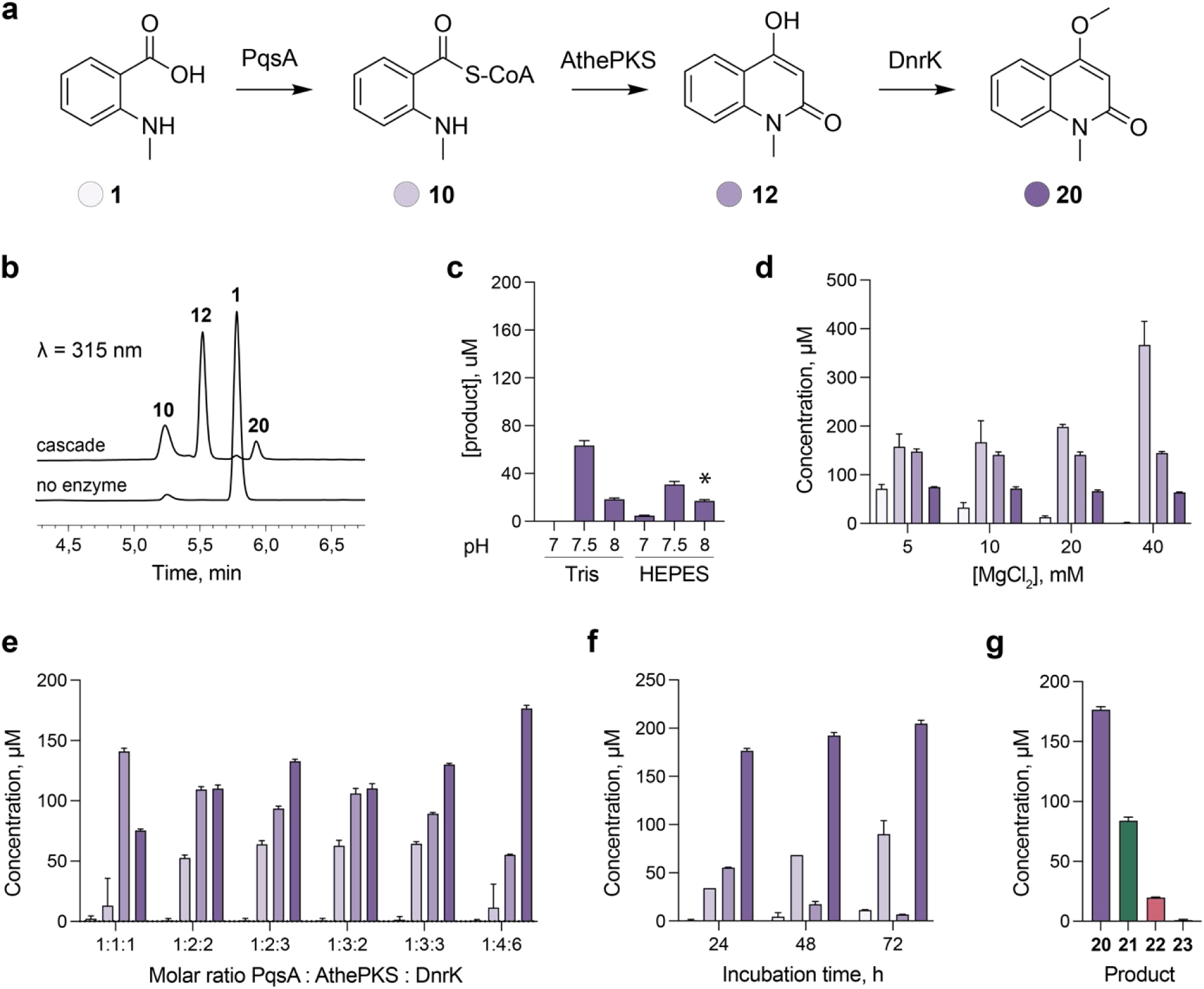
Assembling a one-pot biocatalytic cascade for the production of **20**. a) The artificial pathway towards **20** from **1** featuring three enzymes – PqsA, AthePKS and DnrK. b) Analysis of the reaction products of the cascade under initial conditions and “no enzyme” control by High Performance Liquid Chromatography (HPLC); detection at λ = 315 nm. Initial conditions included 100 mM HEPES-NaOH pH 8, 500 µM **1**, 5 mM MgCl_2_, 2 mM ATP, 250 µM CoA, 500 µM **9**, 2 mM SAM, 1 mol% PqsA, 1 mol% AthePKS, 1 mol% DnrK relative to **1**. The cascade was further optimized by comparing the concentration of **20** and intermediates at c) different buffer pH (asterisk indicates the starting conditions); d) varying concentrations of MgCl_2_ in the reaction mix; e) varying molar ratios of the enzymes in the reaction mix. f) Concentrations of cascade intermediates upon prolonged incubation at optimised conditions (500 µM **1**, 5 mM MgCl2, 2 mM ATP, 62.5 µM CoA, 500 µM **9**, 2 mM SAM, 1 mol% PqsA, 4 mol% AthePKS, 6 mol% DnrK in 100 mM Tris-HCl pH 7.5; 37 °C). g) Cascade-mediated production of 4-methoxylated heterocycles **20-23**. The data are represented as mean ± SD; n = 3.

We noticed that the final product **20** accounts for 73 ± 5% of the total cascade intermediates already after 24 h, despite its concentration corresponding to only 35% of the theoretical maximum. To assess whether a side/back reaction was taking place, we incubated **12** and **20** in the reaction mix with or without the enzymes for up to 72 h (Supplementary Figure 9). The final product **20** was relatively stable in the reaction mix, with 90% of it still detectable after 72 h at 37 °C. However, the concentration of the quinolone intermediate **12** decreased by 20% after 24 h and by 40% after 72 h. The disappearance of **12** was accompanied by the accumulation of two new peaks with *m/z* 363 and *m/z* 453 (positive ion mode), which we had also detected in the cascade reactions (Supplementary Figure 10). The tandem mass spectrometry (MS/MS) fragmentation and UV-Vis absorbance spectra of the side product with *m/z* 363 were similar to those of zanthobisquinolone – a dimer of **12** previously isolated from *Zanthoxylum simulans*^32^. Given its increased abundance in the “no enzyme” control, its formation is likely non-enzymatic. The formation of the other side product, on the other hand, was time-dependent and increased in the mix containing enzymes. Thus, while its identity is elusive, its formation could be suppressed by increasing the efficiency of the methylation step.

Lastly, we evaluated the performance of the optimised cascade for the production of other 4- methoxylated heterocycles **21**-**23** with **2**, **6** and **8** as the starting substrates. We observed the formation of the expected products with all substrates, although 4-methoxythiocoumarin **23** was only present in trace amounts (Figure 4g).

### *In vivo* implementation of the artificial pathway

The use of expensive cofactors (ATP, CoA and SAM) was one of the cost- and scale-limiting factors in the implementation of the cascade towards **20**. The cofactor requirement could be alleviated by engineering the pathway in a microbial cell factory, where the cofactors can be produced and recycled *in situ*. Furthermore, the starting substrate **1** can in principle be derived from anthranilic acid, a common primary metabolite in microbes. Therefore, we endeavoured to implement the cascade in a microbial host and compare the efficiency of the two approaches for the production of **20**. We selected *E. coli* because of the ease of manipulation and because the antimicrobial activity of **20** reportedly does not extend to Gram-negative bacteria^24^.

We cloned PqsA and AthePKS into pETDuet-1 and DnrK into pRSFDuet-1 and co-transformed the resulting plasmids into *E. coli* BL21(DE3), generating strain s1. After the initial fermentation with fed substrate **1** (final concentration 200 µM), we observed a small peak of the quinolone intermediate **12**, but only trace amounts of the final product **20** (Figure 5a). We then explored different plasmid configurations by cloning PqsA and AthePKS into the high copy number plasmid pRSFDuet-1 and DnrK into the medium copy number plasmid pCDFDuet-1, generating strain s2. We also generated a plasmid combination inspired by the enzyme ratio that worked best in the cascade setting: PqsA in pCDFDuet-1, and DnrK and AthePKS in pRSFDuet-1, generating strain s3. Counterintuitively, s3 provided only a marginal increase in the titre of **20** compared to s1. In s2, however, the production of **20** increased to quantifiable levels and averaged at 2 µM.

**Figure 5.**
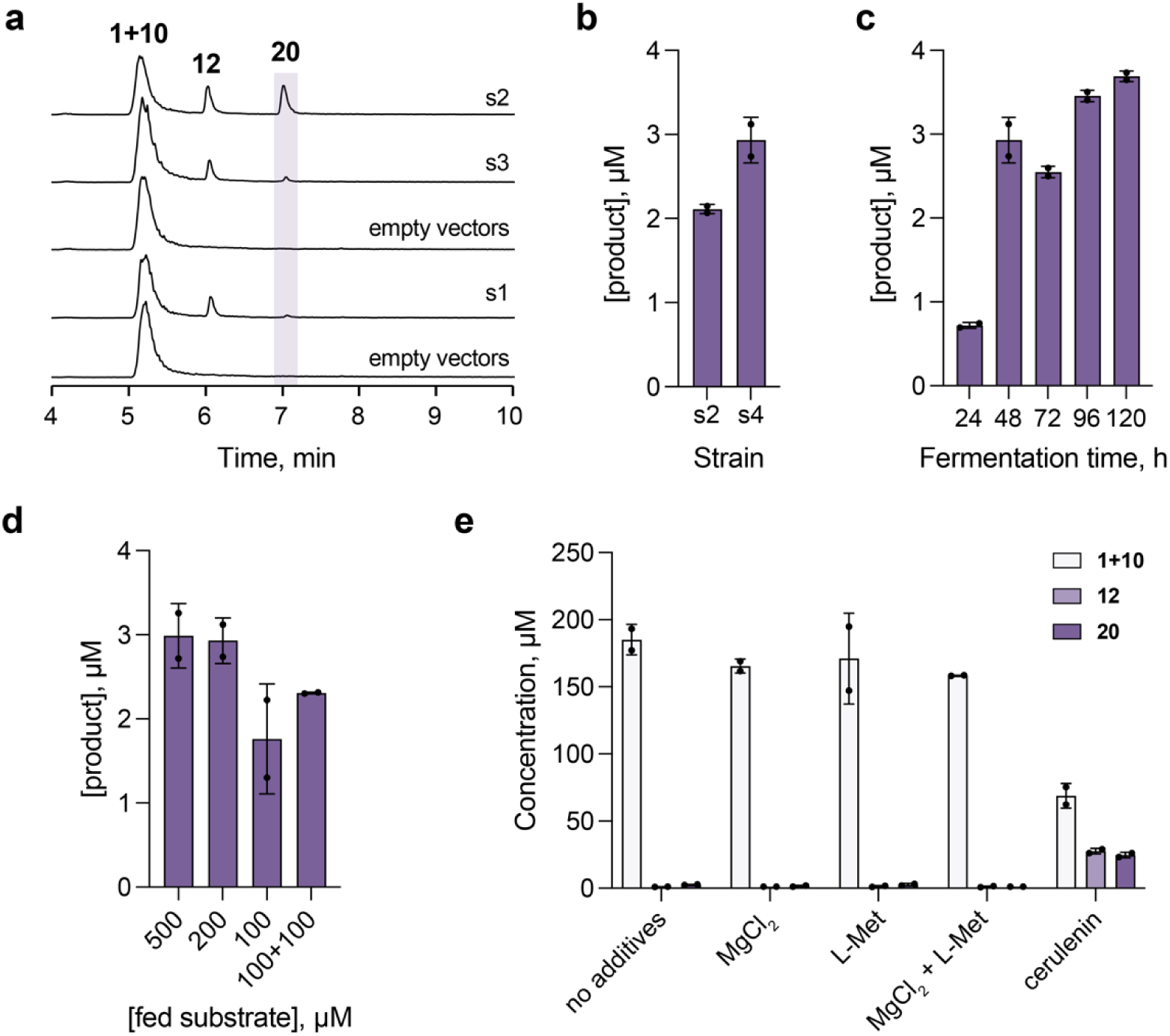
Engineering the biosynthetic pathway towards **20** in *E. coli*. a) Extracted ion chromatograms of the pathway intermediates across engineered *E. coli* strains. s1: PqsA and AthePKS in pETDuet-1, DnrK in pRSFDuet-1; s2: PqsA and AthePKS in pRSFDuet-1, DnrK in pCDFDuet-1; s3: PqsA in pCDFDuet-1, DnrK and AthePKS in pRSFDuet-1. b) Production of **20** in *E. coli* BL21(DE3) (s2) and MG1655(DE3) (s4). c) Time course of **20** accumulation in *E. coli* MG1655(DE3). d) Influence of substrate concentration on the titre of **20**. In “100+100” samples, 100 µM of **1** were added to the culture once every 24 h. e) Effect of additives (10 mM MgCl_2_ and/or 2 mM L-methionine; 20 µg/mL cerulenin) on the concentration of **20** and pathway intermediates **1** and **12**.

We then co-transformed the best combination of plasmids into *E. coli* MG1655(DE3), a K-12 derivative widely used in research due to its well-characterized genome and lack of certain mobile genetic elements like F plasmids or lambda phage^33^. The obtained strain s4 afforded 3 µM product after 48 h of fermentation and was used for all following experiments (Figure 5b). A time-course fermentation experiment suggested that incubation beyond 48 h provides little increase in the titre of **20** (Figure 5c). Feeding higher concentration of substrate or adding it in batches also did not improve the production of **20** (Figure 5d). Lastly, we explored several additives aimed at increasing precursor supply for the T3PKS and OMT reactions (Figure 5e). The addition of 10 mM MgCl_2_ and/or 2 mM L-methionine, the precursor of the methyl donor SAM, did not improve the titre of **20** compared to the “no additives” control. However, the addition of cerulenin, which inhibits fatty acid synthesis and increases the intracellular pool of malonyl-CoA^34^, increased the titre of **20** to ∼25 µM (12% conversion).

Thus, we successfully engineered a biosynthetic pathway to produce **20** in *E. coli* for the first time. Through copy number optimisation and the addition of cerulenin, we were able to increase the product titre from barely detectable levels to ∼25 µM. Increasing the availability of malonyl-CoA was essential to improve the production of **20**, but the cost of cerulenin would make its use in large-scale fermentations impractical. Alternative strategies could be employed to increase the availability of malonyl-CoA, such as the recently developed artificial non-carboxylative malonyl-CoA pathway^35^ and several genetically encoded strategies to arrest fatty acid synthesis^36,37^. However, even with the addition of cerulenin, most of the substrate was not converted to **12**, suggesting that the T3PKS is a major bottleneck in the pathway. Thus, we believe that increasing the catalytic efficiency of AthePKS with **10** is an essential step to improve the yield of the cascade. Therefore, we turned to structural characterisation of AthePKS to identify hotspots for future engineering efforts.

### Structural characterization of AthePKS

We determined the crystal structure of apo-AthePKS at 1.9 Å (Supplementary Table 1). A canonical homodimer with conserved αβαβα thiolase fold was present in the asymmetric unit, which agrees with the expectation that T3PKS form homodimers in solution (Figure 6a). The electron density at the catalytic cysteine (C155) is consistent with its oxidation into sulfinic acid – a phenomenon observed in most T3PKS structures determined to date and thought to reflect the intrinsic redox potential and reactivity of this residue^38^. Structurally, AthePKS shares most similarity with ORAS from *Neurospora crassa* (PDB: 3EUQ, Z-score 57.9, RMSD 1.5 Å)^39^, followed by CsyB from *Aspergillus oryzae* (PDB: 3WXZ, Z-score 54.7, RMSD 2.1 Å)^40^ – the only fungal T3PKSs crystallised to date. Like in ORAS, the active site cavity of AthePKS is connected to a long tunnel lined by hydrophobic residues (Figure 6b). The presence of this tunnel suggests that fatty acyl substrates are likely the native substrates of AthePKS, and its activity with **10** is promiscuous rather than physiological. Furthermore, structural alignment yielded hits to the amoebal Steely1^41^ and chalcone synthases from ancient vascular plants^38,42,43^, as well as several divergent T3PKSs from flowering plants (Supplementary Table 2).

**Figure 6.**
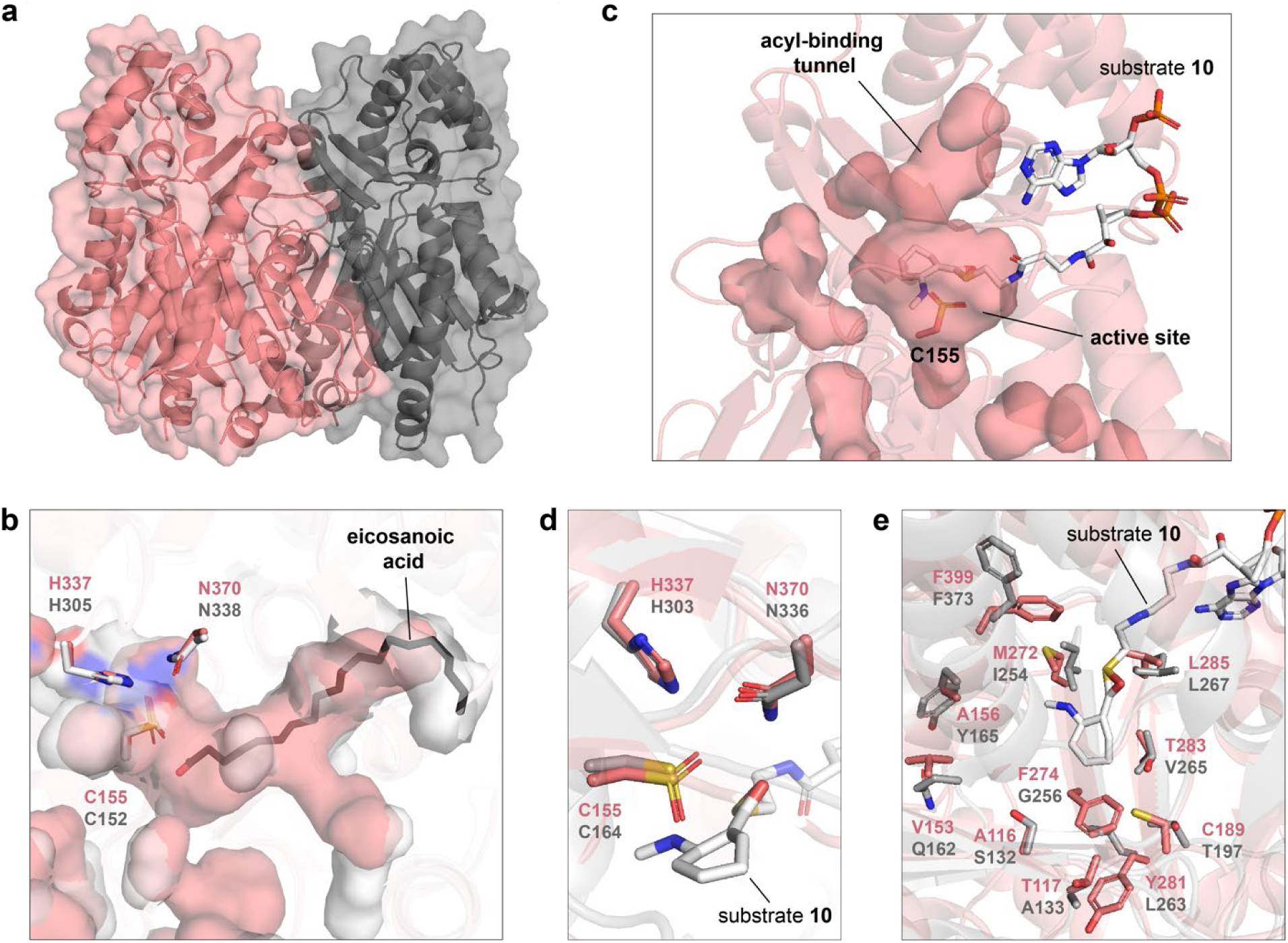
Structural characterization of AthePKS. a) Cartoon representation of the AthePKS dimer (PDB: 9HRB). b) Comparison of the hydrophobic tunnel connected to the active site in ORAS (carbon atoms – grey; PDB: 3EUT) and AthePKS (carbon atoms – pink). The catalytic triad residues and eicosanoic acid from 3EUT are shown as sticks (other atoms: nitrogen – blue, oxygen – red, sulphur – yellow). c) Superimposition of substrate **10** from the structure of AmQNS (white; PDB: 7CCT) with the structure of AthePKS. d) Superimposition of the catalytic triad residues in AmQNS (carbon atoms – grey) and AthePKS (carbon atoms – pink). e) Active site architecture of AthePKS compared to AmQNS. Catalytic triad and several consensus residues (L214, V271, S338, P375; residue numbering from AmQNS) are hidden for visibility.

Despite numerous attempts, co-crystallisation and soaking of AthePKS with **10**, **12** or **16** failed to yield ligand-bound crystals. Instead, we compared the active site of AthePKS to that of the quinolone synthase AmQNS from *Aegle marmelos*^44^ (PDB: 7CCT), which is complexed with substrate **10** (Figure 6c). We observed an almost ideal superimposition of the catalytic triad residues (C164, H303 and N336 in AmQNS, Figure 6d). Position 265, which is occupied by a “gatekeeper” phenylalanine in most T3PKSs and is critical for substrate recognition^45^, is a valine in AmQNS and a threonine in AthePKS (T283, Figure 6e). A smaller residue in this position is associated with the broadening of the active site entrance, which facilitates the access for the bulky substrate **10** in quinolone synthases and related enzymes acridone synthases^14,15,46^.

S132 and A133 in AmQNS are substituted with A116 and T117 in AthePKS. These positions were previously shown to control starter substrate specificity and ultimately the preferred cyclisation mechanism of AmQNS and acridone synthase from *Ruta graveolens*^14,46^. In addition, bulky substitutions I254M and G256F in AthePKS are limiting the volume of its substrate-binding pocket compared to AmQNS and related enzymes (Supplementary Figure 11). Manipulation of the highlighted residues might modulate the affinity and catalytic efficiency of AthePKS towards substrate **10** for improved production of **20**.

### Conclusions

In summary, we have developed a one-pot biocatalytic platform for the synthesis of three different heteroaromatic scaffolds. Using a series of anthranilic and salicylic acid derivatives, we demonstrated the production of several pharmaceutical intermediates which had not been produced enzymatically before. Even under unoptimised conditions, AthePKS or FerePKS showed moderate to good conversions with substrates **1**, **3**, **6** and **7**. In the case of thiocoumarin **18**, our two-step biocatalytic strategy even offers an advantage compared to the conventional 4- step chemical synthesis^47^. The crystal structure of AthePKS determined in this study may facilitate engineering of the enzyme to further increase its catalytic efficiency with a particular substrate.

Using AthePKS, we developed an artificial enzymatic cascade towards an antimicrobial *Rutaceae* alkaloid **20**, achieving its biosynthesis for the first time. To bridge the gap in its cryptic biosynthetic pathway, we searched the literature for candidate *O*-methyltransferases and identified an enzyme that, apart from the target intermediate, was active on all other heteroaromatic scaffolds. We also demonstrated the production of unnatural 4-methoxylated (thio)coumarins through changing the starter substrate in the cascade.

While this work provides a framework for the cascade biosynthesis of **20** both *in vitro* and in a microbial host, further optimisation is necessary to render these strategies competitive with chemical synthesis. The performance of the cascade is currently superior *in vitro* but we see several options to improve it further. First, increasing the efficiency of *O*-methylation, either through protein engineering or homologue screening, could outrun the formation of side products from the quinolone intermediate. Together with the engineering of the T3PKS, it would also allow to increase the substrate load above 1 mM. Second, the components of the cascade can be used more efficiently. Currently, the first step of the cascade recycles the CoA generated from malonyl-CoA condensation during the T3PKS reaction. This recycling aspect can be taken one step further by introducing a malonate-CoA ligase^48^. *In situ* generation of malonyl-CoA, which is currently the most expensive component of the cascade, would also enable scale-up. Finally, incorporating a SAH degradation/SAM regeneration system would address both the efficiency of methylation and cofactor economy in the cascade^49,50^. Future experimental efforts should focus on incorporating these strategies for efficient enzymatic synthesis of **20** and other pharmaceutically valuable quinolones/(thio)coumarins.

## Materials and methods

### Construction of plasmids

The list of all plasmids used in this study is provided in Supplementary Table 3. Plasmids for the expression of PqsA, AthePKS, FerePKS, GloKOMT and MesMOMT were constructed in our previous studies^18,31^. The genes coding for DnrK, CmOMT2 and CpdC were codon-optimised for expression in *E. coli* and ordered as linear DNA fragments from Twist Bioscience (San Francisco, USA). The genes coding for DnrK, CmOMT2 and CpdC with N-terminal hexahistidine tag were cloned by restriction and ligation (NcoI/HindIII) into pRSFDuet-1 for expression under the T7 promoter. For pathway applications, different combinations of the pathway genes were cloned between pETDuet-1, pRSFDuet-1 and pCDFDuet-1 using restriction and ligation or Gibson assembly. All constructs were verified by Sanger sequencing (Macrogen Europe, Amsterdam). Amino acid sequences of all proteins expressed in this study can be found in Supplementary File 1.

### Protein expression, purification and crystallisation

#### Expression

The expression and purification procedures were the same for PqsA, AthePKS, FerePKS, DnrK, CmOMT2 and MesMOMT. Plasmids harbouring the target genes were transformed into chemically competent *E. coli* BL21(DE3) and maintained on selective LB agar containing the appropriate antibiotic. A starter culture was inoculated from a single colony (5 mL, LB with appropriate antibiotic) and incubated at 37 °C (180 rpm, overnight). The main culture was inoculated from the starter culture (1 : 100) into 400 mL of TB autoinduction medium with appropriate antibiotic (20 g/L tryptone, 24 g/L yeast extract, 4 ml/L glycerol, 17 mM KH_2_PO_4_, 72 mM K_2_HPO_4_, 0.2 % w/v lactose and 0.05 % w/v glucose) and incubated at 37 °C (180 rpm, 2 h), after which the temperature was lowered to 25 °C (180 rpm, overnight).

#### Purification

The cells were harvested by centrifugation (15 min, 3,428 × g) and the pellet was resuspended in 5 volumes of lysis buffer (buffer A including one EDTA-free protease inhibitor tablet (Roche, Basel, Switzerland); buffer A: 50 mM Tris/HCl pH 7.5, 500 mM NaCl, 20 mM imidazole). The cell suspension was lysed by sonication (50 % duty cycle, 7 cycles of 35 s ON/60 s OFF) and cleared by centrifugation for 60 min at 17,000 × g. The supernatant was loaded onto the Ni-NTA affinity matrix (Qiagen, Hilden, Germany) equilibrated with buffer A by gravity flow. The column was washed with 20 column volumes of wash buffer (50 mM Tris/HCl pH 7.5, 500 mM NaCl, 30 mM imidazole) and eluted stepwise with one column volume of buffers B1 to B6 (buffers B1-B6: 50 mM Tris/HCl pH 7.4, 500 mM NaCl, 50 mM imidazole/ 100 mM imidazole/ 150 mM imidazole/ 200 mM imidazole/ 250 mM imidazole or 500 mM imidazole, respectively). The eluates of each step were collected in separate fractions and analysed by SDS-PAGE. Fractions containing target proteins with low background were pooled and transferred into the storage buffer (10 mM HEPES/NaOH pH 7.5, 50 mM NaCl, 2 mM dithiothreitol, 5% (v/v) glycerol) by 3 cycles of concentration (Amicon® Ultra Centrifugal Filter; 3 kDa molecular weight cutoff) and dilution (1:30). For protein crystallisation experiments, AthePKS was additionally subjected to size-exclusion chromatography on a Superdex 75 pg column (GE Healthcare, USA) and eluted into the same storage buffer. The protein concentrations were determined by absorbance at 280 nm (NanoDrop, ThermoFisher Scientific, USA) before the purified enzymes were aliquoted and flash-frozen with liquid nitrogen for storage at −70 °C. SDS-PAGE analysis of the purified proteins is shown in Supplementary Figure 12.

#### Crystallisation

The crystallization of AthePKS was performed using the sitting-drop vapor diffusion method at a temperature of 18 °C. A 1:1 μL mixture of reservoir solution and freshly prepared protein was employed. Initial crystallization screening utilised commercial kits, specifically JCSG+ and Morpheus from Molecular Dimensions Co. Subsequently, optimisations were performed to obtain higher quality crystals. Tetragonal AthePKS crystals were observed at a protein concentration of 10 mg/mL in a crystallization buffer composed of 0.1 M Tris (pH 8.5), 0.03 M MgCl_2_, 0.03 M CaCl_2_, 10% (w/v) PEG8000, and 20% (w/v) PEG200. Crystals were harvested using nylon loops 3-4 days prior to diffraction. The crystals were briefly immersed in cryoprotectant solutions, prepared by supplementing the reservoir solutions with 20-30% (v/v) glycerol, before being flash-cooled in liquid nitrogen.

#### Diffraction data collection, structure determination and refinement

Diffraction data were collected at beamline P11 of the PETRA III facility (DESY, Hamburg, Germany). The collected data were auto-processed using XDSAPP^51^, scaling and merging statistics were subsequently handled with Aimless in the CCP4 suite^52^. The structure of AthePKS was determined by molecular replacement. The monomer of type III polyketide synthase from *Neurospora crassa* (PDB ID: 3E1H)^53^ was utilized as the search model, employing the program MOLREP^54^. Further refinement of the structure was carried out using REFMAC5^55^ and manual adjustments made in Coot^56^. The data collection and refinement statistics are presented in Supplementary Table 1.

The structure of AthePKS was deposited in the PDB under accession code 9HRB.

### Chemical synthesis of reference compounds

For the synthesis of methoxylated heterocycles **20**-**23**, trimethylsilyldiazomethane (2 M in hexane, 1.5 mL, 3 mmol) was added to a solution of the starting material (**12**, **13**, **16** or **18**; 1 mmol) in MeOH (2 mL) and DCM (2 mL) at 0°C under N_2_. The reaction mixture was warmed to RT, stirred for 30 min (controlled by TLC), and then concentrated and purified by MPLC (4 g, 0 - 100% EtOAc/Pentane) to provide the product as an off-white solid. ^1^H NMR spectra of all chemically synthesised compounds can be found in Supplementary Figures 13-16.

### *In vitro* assays

#### PqsA substrate scope analysis

To probe the substrate scope of PqsA, reactions were set up in triplicate with carboxylic acids **1- 8** and a “no substrate” control and contained 2 mM ATP, 2 mM MgCl_2_, 0.5 mM substrate, 0.2 mM CoA and 5 μM PqsA in 100 mM Tris-HCl pH 7.5; reaction volume: 50 µL. Additional negative control reactions were set up under identical conditions lacking ATP. The reactions were incubated at 37 °C for 2 h without shaking and analysed by quantifying the residual-SH groups of coenzyme A in the reaction mixture using 5,5′-dithiobis-(2-nitrobenzoic acid) (DTNB, Ellmann’s reagent) as described previously^57^.

#### T3PKS substrate scope analysis

The substrate scope of AthePKS and FerePKS was assessed in a one-pot continuous cascade with PqsA. The reaction mix consisted of 0.5 mM substrate **1-8**, 20 mM MgCl_2_, 2 mM ATP, 0.25 mM CoA, 0.5 mM malonyl-CoA (**9**) or methylmalonyl-CoA (**11**) in 100 mM Tris-HCl pH 7.5; reaction volume: 30 µL. 5 µM PqsA and 5 µM Athe/FerePKS were added simultaneously. For each substrate, negative control reactions were set up under identical conditions lacking PqsA and the T3PKS. The reactions were incubated at 37 °C for 24 h without shaking, after which an equal volume of acidic methanol (MeOH + 0.1% formic acid) was added to precipitate the proteins. The reactions were centrifuged at 17,000 × *g* for 10 min and analysed by LC-MS and/or HPLC for quantification.

#### Screening of the OMTs using cell-free extracts

To screen for the desired quinolone 4-*O*-methyltransferase activity, the candidate genes were expressed from plasmids c173, c192 and c379-c381 in *E. coli* as described in “Protein expression, purification and crystallisation” (culture volume: 10 mL). 1 mL of each culture was centrifuged at 4500 × g for 10 min, and the pellets were incubated at –70 °C for 45 min. Afterwards, the cells were thawed and resuspended in 200 µL of lysis buffer (25 mM Tris-HCl pH 7.5, 1 mg/mL lysozyme and 1:10000 benzonase (Novagen)) and incubated at 30 °C without shaking for 1 h. Following the incubation, 600 µL of 25 mM Tris-HCl pH 7.5 were added, and the insoluble cell fraction was removed by centrifugation at 17,000 × g for 15 min. 20 µL of each cell-free lysate were then used for OMT assays together with 200 µM substrate **12**, 5 mM MgCl_2_ and 1 mM SAM in 25 mM Tris-HCl pH 7.5; final volume: 100 µL. The reactions were incubated at 30 °C for 24 h, quenched with an equal volume of acidic methanol to precipitate the proteins, centrifuged at 17,000 × *g* for 10 min and analysed by LC-MS.

#### *In vitro* OMT assays

Substrate scope of the purified DnrK, CmOMT2 and MesMOMT was assessed in triplicate by incubating 0.5 mM substrate **12**, **13**, **16** or **18**, 2 mM SAM and 5 µM enzyme in 100 mM HEPES-NaOH pH 8; reaction volume: 30 µL. To assess the influence of Mg^2+^ on the activity of the OMTs, an identical set of reactions was prepared with the addition of 5 mM MgCl_2_. For each substrate, negative control reactions were set up under identical conditions lacking the OMT. The reactions were incubated at 37 °C for 24 h, quenched with an equal volume of acidic methanol to precipitate the proteins, centrifuged at 17,000 × *g* for 10 min and analysed by LC-MS.

#### One-pot three-step biocatalytic cascade

The initial conditions for the three-step cascade included 0.5 mM substrate **1**, 5 mM MgCl_2_, 2 mM ATP, 0.25 mM CoA, 0.5 mM malonyl-CoA and 2 mM SAM in 100 mM HEPES-NaOH pH 8; reaction volume: 30 µL. PqsA, AthePKS and DnrK (1 mol% or 5 µM each) were added simultaneously, and the reactions were incubated at 37 °C for 24 h. For pH and buffer optimisation, HEPES-NaOH pH 8 was substituted for HEPES-NaOH (pH 7 and pH 7.5) or Tris-HCl (pH 7, pH 7.5, pH 8). To assess the effect of varying Mg^2+^ concentration, MgCl_2_ was added at concentrations of 5 mM, 10 mM, 20 mM and 40 mM. To assess the effect of lowering the starting concentration of CoA, reactions were set up with CoA to substrate ratios of 1:1, 1:2, 1:4, 1:8, 1:16 and 1:32. At each stage of the optimisation, negative control reactions were set up under identical conditions lacking the cascade enzymes. For time-course experiments, the reactions were incubated for up to 72 h and samples were taken every 24 h. The optimised conditions included 0.5 mM substrate **1**, 5 mM MgCl_2_, 2 mM ATP, 62.5 µM CoA, 0.5 mM malonyl-CoA and 2 mM SAM in 100 mM Tris-HCl pH 7.5. Cascade reactions with substrates **2**, **6** and **8** were set up under the optimised conditions and incubated for 24 h. All reactions were quenched with an equal volume of acidic methanol to precipitate the proteins, centrifuged at 17,000 × *g* for 10 min and analysed by LC-MS.

### Fermentation

Strains harbouring the quinolone pathway genes were generated by transforming the appropriate combinations of plasmids into chemically competent^58^ *E. coli* BL21(DE3) or electrocompetent^59^ *E. coli* MG1655(DE3). The list of all strains used in the fermentation experiments is provided in Supplementary Table 4. Starter cultures were prepared from two individual colonies of the final strains in 5 mL LB supplemented with appropriate antibiotics. The cultures were incubated overnight at 37°C with agitation, and 100 µL thereof were used to inoculate the main cultures (10 mL TB-autoinduction with antibiotics; sterile glass tubes). The substrate was added after 3 h of growth at 37°C, 200 rpm (OD_600_ ∼ 0.6). These cultures were incubated at 30°C, 200 rpm for 48 h. Samples of 667 μL were taken after 48 h and extracted with the double volume (1334 µL) of acidic methanol (MeOH + 0.1% formic acid). The extracts were dried, resuspended in 25 µL of acidic methanol and analysed by LC-MS. For the time-course experiment, the cultures were incubated for 120 h, and samples of 667 µL were taken every 24 h. When required, 10 mM MgCl_2_, 2 mM L-methionine or 20 µg/mL cerulenin were added to the cultures at the time of substrate addition.

### Analysis and quantification of target compounds

The detection of *in vitro* reaction and fermentation products was performed using a low-resolution Waters Acquity Arc UHPLC-MS system equipped with a 2998 PDA detector and a QDa single-quadrupole mass detector. The samples were separated over a Waters XBridge BEH C18 3.5 μm 2.1×50 mm column at 40 °C with a concentration gradient (solvent A: water +0.1 % formic acid, and solvent B: acetonitrile + 0.1 % formic acid) at a flow rate of 0.5 mL/min (2 μL injections). The following gradient was used: 5 % B for 2 min, 5–90 % B over 3 min; 90 % B for 2 min; 5 % B for 3 min. For *E. coli* fermentation extracts, a longer gradient was employed: 5% B for 2 min, 2– 50% B over 8 min, 50–90% B over 5 min, 90% B for 2 min, 5% B for 3 min. MS analysis was carried out in both positive and negative ion modes with the following parameters: probe temperature of 600 °C; capillary voltage of 1.0 kV; cone voltage of 15 V; scan range 100-1250 *m/z*. The acquired data were analysed using the proprietary software MassLynx.

The identity of the reaction products was further confirmed by high-resolution and MS/MS analyses using a Shimadzu LC20-XR system (Shimadzu Benelux, Den Bosch, The Netherlands) coupled to a Q Exactive Plus mass spectrometer (Thermo Fisher Scientific, USA). The samples were separated over a Waters XBridge BEH C18 reversed-phase column at 50 °C with a concentration gradient (solvent A: water + 0.1 % formic acid, and solvent B: acetonitrile + 0.1 % formic acid) at a flow rate of 0.5 mL/min (2 μL injections). The following gradient was used: 5 % B for 2 min, 5–90 % B over 3 min; 90 % B for 2 min; 5 % B for 3 min. MS and MS/MS analyses were performed with electrospray ionization (ESI) in positive or negative ion mode at a spray voltage of 3.5, and sheath and auxiliary gas flow set at 48 and 11, respectively. The ion transfer tube temperature was 255 °C. Spectra were acquired in data-dependent mode with a survey scan at m/z 80 − 1200 at a resolution of 70,000 followed by MS/MS fragmentation of the top 5 precursor ions at a resolution of 17,500. A stepped collision energy of 30-40-55 was used for fragmentation, and fragmented precursor ions were dynamically excluded for 10 s. The acquired data were analysed using MZmine 4.0.3^60^. Spectral data of all enzymatic products and reference compounds can be found in Supplementary Figures 17-28.

Quantification of the reaction products was performed using reversed-phase HPLC (instrument: Shimadzu LC-20AD; autosampler: SIL-20AC, T=15 °C, 10 μL injection; flow rate: 1.2 mL/min; column: NUCLEOSHELL RP18 90 Å, 100 x 4.6 mm, 2.7 μm, T=40 °C; detector: SPD-20A, λ=355 nm (N-methylanthranilic acid and N-methylanthraniloyl-CoA) and λ=315 nm (4-hydroxy-1-methyl-1-quinolone, 4-methoxy-1-methyl-2-quinolone and all other T3PKS products); solvent A: water with 0.1 % FA, solvent B: ACN with 0.1 % FA; gradient: 10 % B for 2 min; 10–95 % B over 3 min; 95 % B for 2 min; 95–10 % B over 3 min). The peak areas were integrated and converted to concentrations in μM based on calibration curves with the authentic standards (Supplementary Figures 29-31).

## Author contributions

N.S. and K.H. conceived the study. N.S. performed all biochemical experiments and analysed the data. A.O. performed the chemical synthesis and compound characterisation under supervision of S.S. L.Z. and M.G. determined and refined the crystal structure. N.S. wrote the manuscript with contributions from L.Z., A.O., M.G., S.S. and guidance from K.H. All authors read and approved the final version of the manuscript.

## Notes

The authors declare no competing financial interest.

## Acknowledgements

The authors acknowledge DESY (Hamburg, Germany) for the provision of experimental facilities at PETRA III. The authors are grateful to the Interfaculty Mass Spectrometry Center of the University of Groningen and the University Medical Center Groningen for their services in high resolution tandem mass spectrometry.

## Abbreviations

ATP: adenosine triphosphate
CoA: coenzyme A
HPLC: high performance liquid chromatography
LC-MS: liquid chromatography-coupled mass spectrometry
MS/MS: tandem mass spectrometry
NP: natural product
OMT: O-methyltransferase
SAH: *S*- adenosylhomocysteine
SAM: *S*-adenosylmethionine
T3PKS: type III polyketide synthase

