## Supplementary Figures and Tables for "A biocatalytic platform for the production of substituted 2-quinolones and (thio)coumarins"

### Supplementary Tables

**Supplementary Table 1.** Diffraction data collection, structure determination, and refinement statistics.

| **Data set** | | **AthePKS** |
|---|---|---|
| PDB entry | | 9HRB |
| Diffraction source | | beamline P11, DESY |
| Wavelength (Å) | | 1.0332 |
| Temperature (K) | | 100 |
| Detector | | EIGER2 2X 16M |
| Rotation range per image (°) | | 0.1 |
| Total rotation range (°) | | 180 |
| Crystal-to-detector distance (mm) | | 198.874 |
| Space group | | P21 21 21 |
| Unit cell parameters | a,b,c(Å) | a = 71.42 |
| | | b = 89.99 |
| | | c = 141.28 |
| | α, β, γ(°) | α = 90.00 |
| | | β = 90.00 |
| | | γ = 90.00 |
| Resolution, Å | | 47.08-1.90(1.94-1.90) |
| Number of observations | | 495336(31366) |
| Completeness, % | | 99.9(100.0) |
| Multiplicity | | 6.8(7.1) |
| Rmerge | | 0.084(1.167) |
| Average I/σ,I | | 13.8(1.7) |
| CC(1/2) | | 0.999(0.772) |
| Wilson B-factor, Å$^2$ | | 29.89 |
| **Refinement statistics** | | |
| No. reflections, all/free | | 72274/3657 |
| Rfactor | | 0.17 |
| Rfree | | 0.205 |
| No. protein atoms | | 5850 |
| No. ligand atoms | | - |
| No. water atoms | | 228 |
| **Average B-factors, Å$^2$** | | |
| Protein | | 37.6 |
| Ligands | | - |
| Water | | 36.96 |
| **RMSD from ideal values** | | |
| Bond lengths, Å | | 0.0083 |
| Bond angles, ° | | 1.757 |
| **Ramachandran plot** | | |
| Favoured, % | | 98.56% |
| Allowed, % | | 1.44% |
| Disallowed, % | | 0.00% |

**Supplementary Table 2.** Structural alignment of AthePKS with known T3PKSs from PDB with the Dali server.

| # | PDB ID & chain | Z score | rmsd | lali | nres | %id | Ligands | Protein name | Donor organism |
|---|---|---|---|---|---|---|---|---|---|
| 1 | 3euq-B | 57.9 | 1.5 | 371 | 379 | 43 | Hexaethylene glycol | ORAS | *Neurospora crassa* (fungus) |
| 2 | 3wxz-A | 54.7 | 2.1 | 372 | 377 | 32 | / | CsyB | *Aspergillus oryzae* (fungus) |
| 3 | 2h84-A | 48.1 | 2.2 | 357 | 363 | 27 | Hexaethylene glycol | Steely1 | *Dictyostelium discoideum* AX4 (amoeba) |
| 4 | 3awj-A | 47.9 | 2.1 | 356 | 380 | 27 | Coenzyme A; sulfate ion | HsPKS1 | *Huperzia serrata* (moss) |
| 5 | 6dx7-C | 47.7 | 2.0 | 355 | 387 | 27 | / | PpCHS | *Physcomitrium patens* (moss) |
| 6 | 7w6g-D | 47.7 | 2.1 | 356 | 383 | 27 | Lauroyl-CoA | TKS-L190G mutant | *Cannabis sativa* (plant) |
| 7 | 2d3m-B | 47.7 | 2.2 | 356 | 395 | 23 | Coenzyme A | PCS | *Aloe arborescens* (plant) |
| 8 | 7yj9-A | 47.5 | 2.1 | 358 | 389 | 26 | Coenzyme A; naringenin | ScCHS1 | *Odontosoria chusana* (fern) |
| 9 | 6dxf-B | 47.4 | 2.2 | 360 | 379 | 25 | / | SmCHS | *Selaginella moellendorffii* (moss) |
| 10 | 1u0u-A | 47.4 | 2.1 | 356 | 389 | 24 | / | STS | *Pinus sylvestris* (plant) |
| 11 | 7cbf-A | 47.4 | 2.1 | 355 | 383 | 24 | Iodide ion; glycerol; imidazole | BPS | *Garcinia mangostana* (plant) |
| 12 | 5wc4-B | 47.3 | 2.2 | 357 | 378 | 25 | Benzoyl-CoA | MdBIS3 | *Malus domestica* (plant) |
| 13 | 6j1m-A | 47.3 | 2.1 | 355 | 381 | 26 | 3-oxopentanedioic acid | AaPKS2 | *Anisodus acutangulus* (plant) |

**Supplementary Table 3.** List of strains and plasmids used in this study

| **Strain** | **Genotype** | **Source** |
|---|---|---|
| *E. coli* DH5α | F– φ80lacZΔ M15 Δ (*lacZYA-argF*) *U169 recA1 endA1 hsdR17* (rK– mK+) *phoA supE44* λ- *thi–1 gyrA96 relA1* | Invitrogen |
| *E. coli* BL21(DE3) | F–ompT hsdSB (rB–, mB–) gal dcm (DE3) | Invitrogen |
| *E. coli* MG1655(DE3) | K12 F- lambda- *ilvG- rfb*-50 *rph*-1 | [1] |

| **ID** | **Backbone** | **Enzyme encoded** | | **Source** |
|---|---|---|---|---|
| | | **MCS1** | **MCS2** | |
| c173 | pET21b | MesMOMT | / | [2] |
| c192 | pET21b | GloKOMT | / | [2] |
| c266 | pET28a | AthePKS | / | [3] |
| c272 | pET28a | FerePKS | / | [3] |
| c288 | pET28a | PqsA | / | [3] |
| c325 | pETDuet | PqsA | AthePKS | this study |
| c379 | pRSFDuet1 | CmOMT2 | / | this study |
| c380 | pRSFDuet1 | CpdC | / | this study |
| c381 | pRSFDuet1 | DnrK | / | this study |
| c382 | pCDFDuet1 | DnrK | / | this study |
| c383 | pRSFDuet1 | PqsA | AthePKS | this study |
| c387 | pCDFDuet1 | PqsA | / | this study |
| c388 | pRSFDuet1 | DnrK | AthePKS | this study |

* MCS1 and 2, multiple cloning site (each with its own T7 promoter and terminator); MesMOMT, OMT from *Mesorhizobium muleiense*; GloKOMT, OMT from *Gloeocapsa* sp. PCC 7428; AthePKS, T3PKS from *Aspergillus thesauricus* IBT 34227; FerePKS, T3PKS from *Fonsecaea erecta* CBS 125763; PqsA, anthranilate-CoA ligase from *Pseudomonas aeruginosa*; CmOMT2, OMT from *Cnidium monnieri*; CpdD, OMT from *Aspergillus californicus*; DnrK, carminomycin OMT from *Streptomyces peucetius*; all codon-optimized for expression in *E. coli*.

**Supplementary Table 4.** List of *E. coli* strains used in fermentation experiments in this study.

| **ID** | **Host strain** | **Construct ID** | **Enzymes expressed** | **Source** |
|---|---|---|---|---|
| s1 | BL21(DE3) | c325, c381 | PqsA, AthePKS, DnrK | this study |
| s2 | BL21(DE3) | c382, c383 | PqsA, AthePKS, DnrK | this study |
| s3 | BL21(DE3) | c387, c388 | PqsA, AthePKS, DnrK | this study |
| s4 | MG1655(DE3) | c382, c383 | PqsA, AthePKS, DnrK | this study |

#### Supplementary Figures

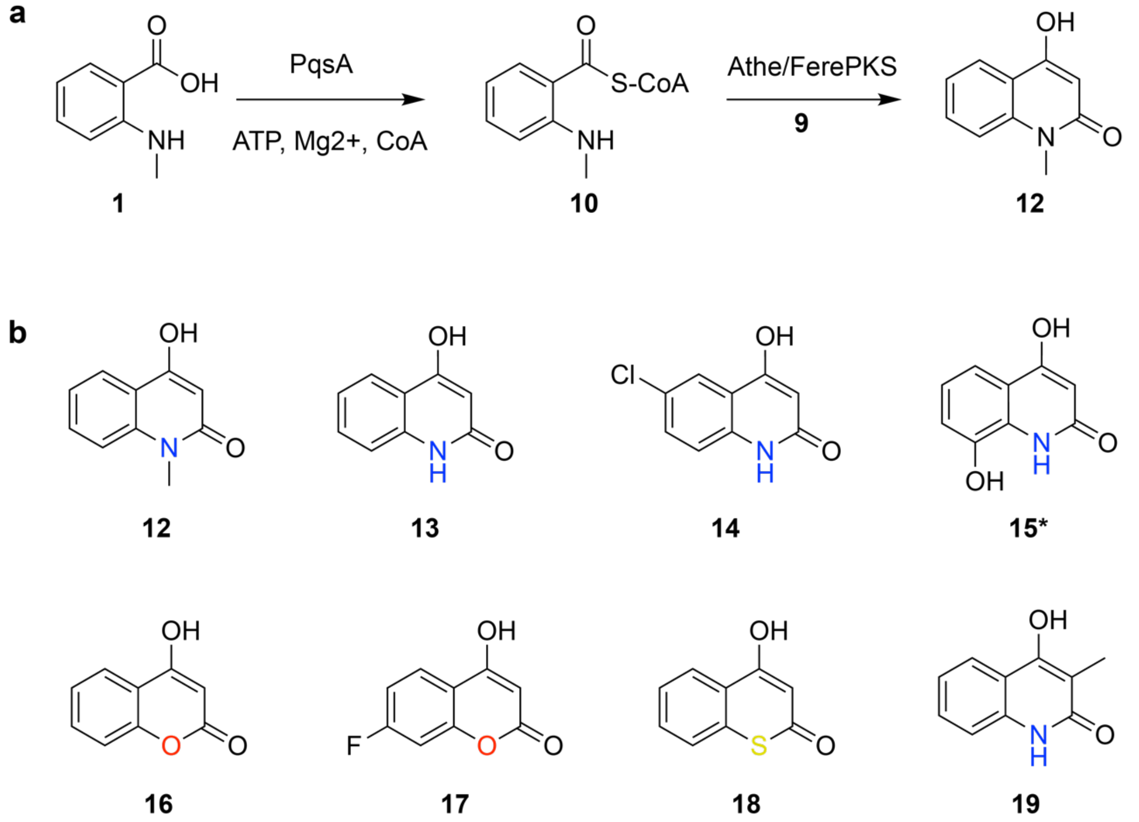

**Supplementary Figure 1.** *In vitro* product scope of the cascade with PqsA and Athe/FerePKS. a) A two-step cascade from **1** to **12**. b) Products **12-19** detected in the cascade reactions. Asterisk indicates the putative product.

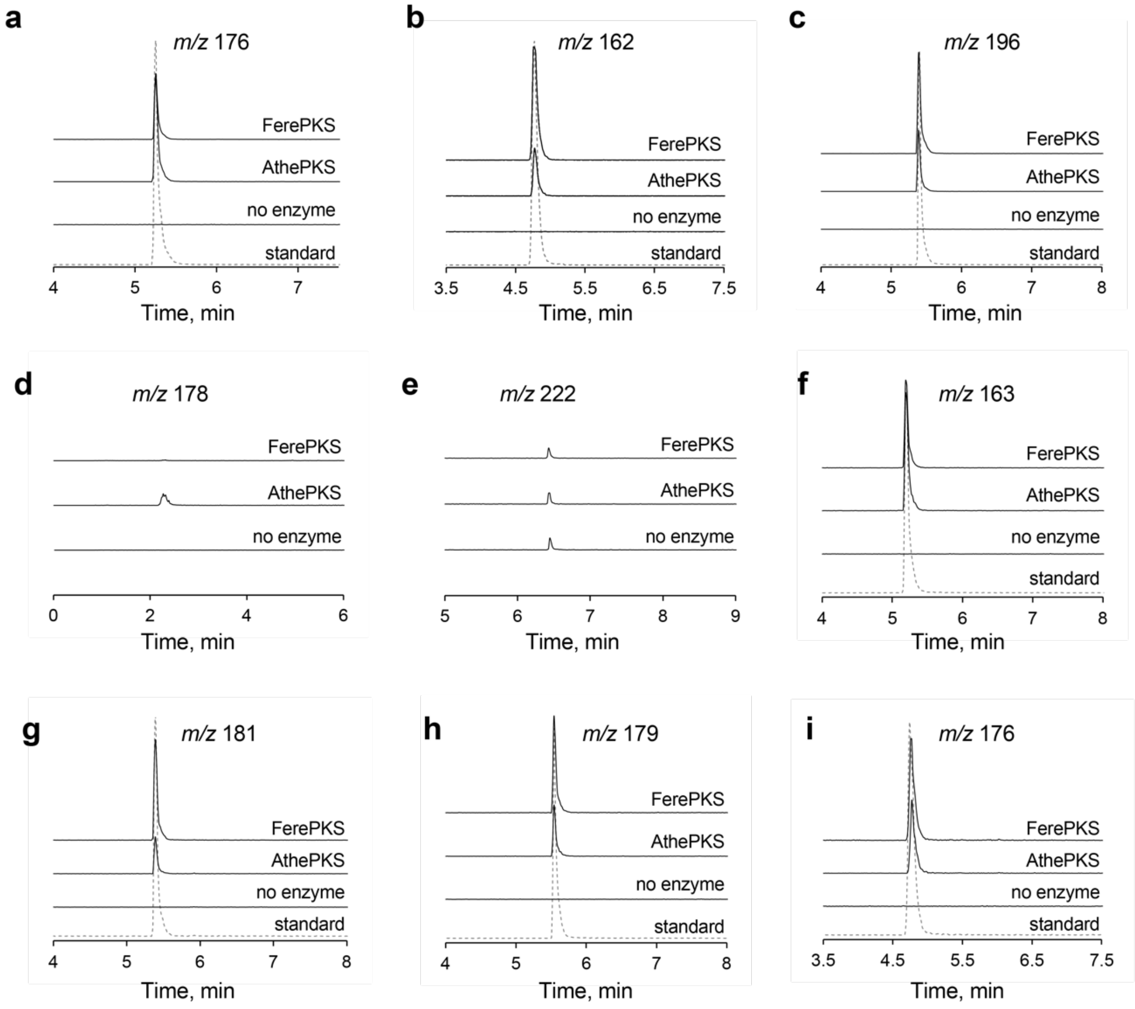

**Supplementary Figure 2.** Extracted ion chromatograms in low-resolution LC-MS of reaction products of Athe/FerePKS with substrates **1-8** compared to the "no enzyme" control and reference compounds. The reactions were incubated as a one-pot continuous cascade with PqsA and malonyl-CoA (**9**) if not stated otherwise. a) Substrate **1**; b) substrate **2**; c) substrate **3**; d) substrate **4**; e) substrate **5**; f) substrate **6**; g) substrate **7**; h) substrate **8**; i) substrate **2** + **11** (methylmalonyl-CoA). In e), an unspecific peak with *m/z* 222 that is present across samples with multiple substrates is shown. It is thus unlikely to be the quinolone product from **5**.

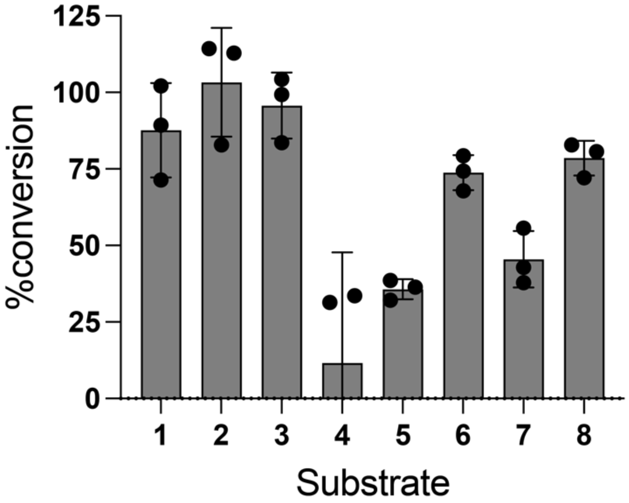

**Supplementary Figure 3.** CoA ligation reactions with PqsA and substrates **1-8**. Conversion was calculated using DTNB assay based on the residual concentration of CoA in the reaction mix as described in the Methods section. The data are represented as mean ± SD; n=3.

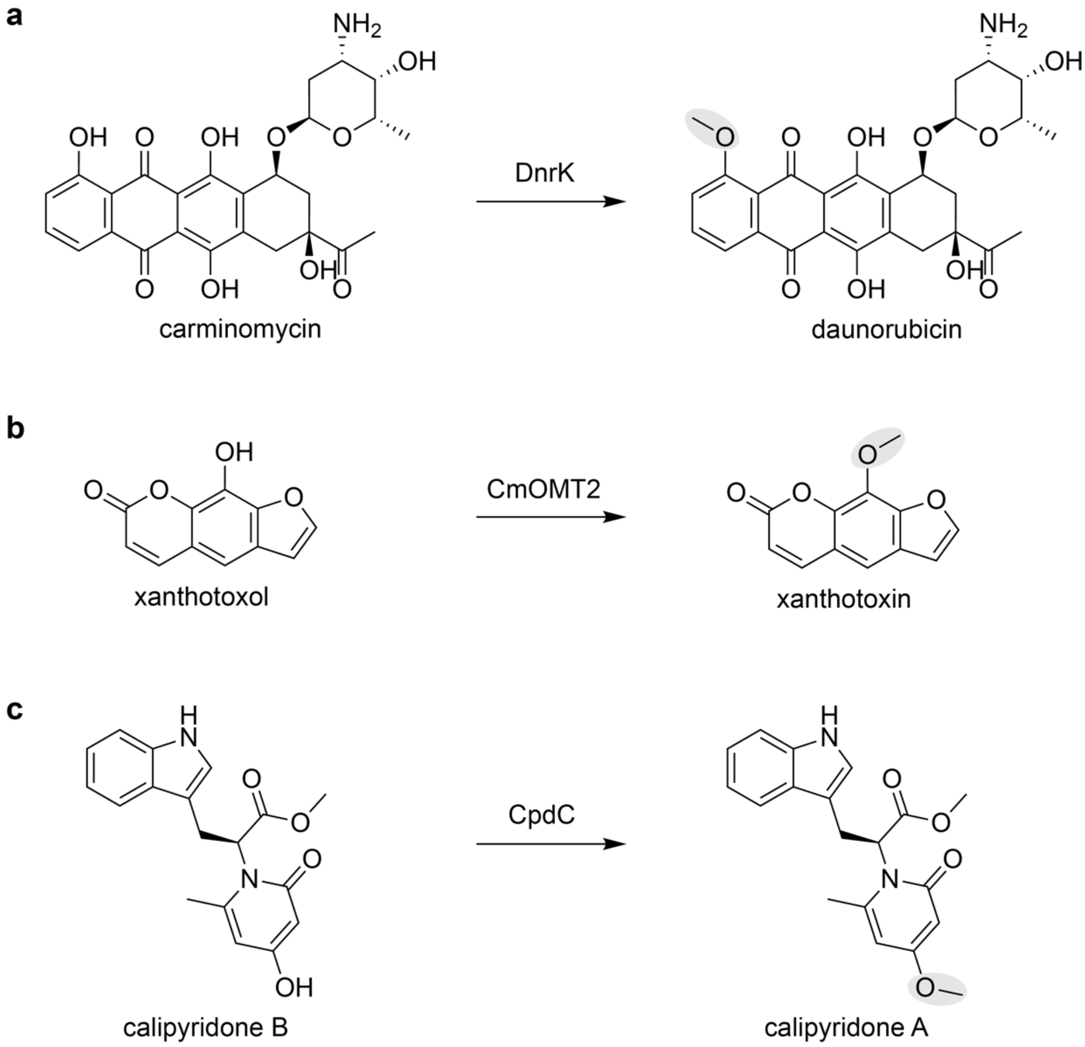

**Supplementary Figure 4.** Native reactions catalysed by a) DnrK, b) CmOMT2 and c) CpdC.

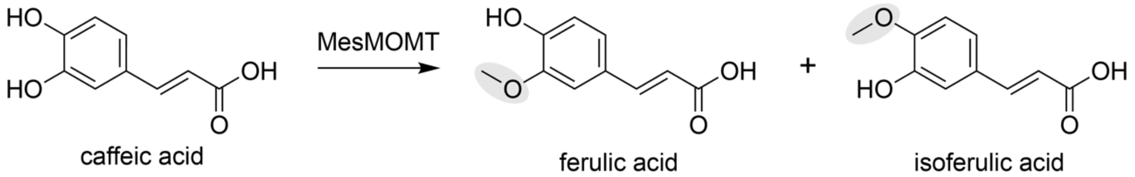

**Supplementary Figure 5.** Scheme of caffeic acid methylation by MesMOMT. GloKOMT was not active on catechol-like substrates in [2].

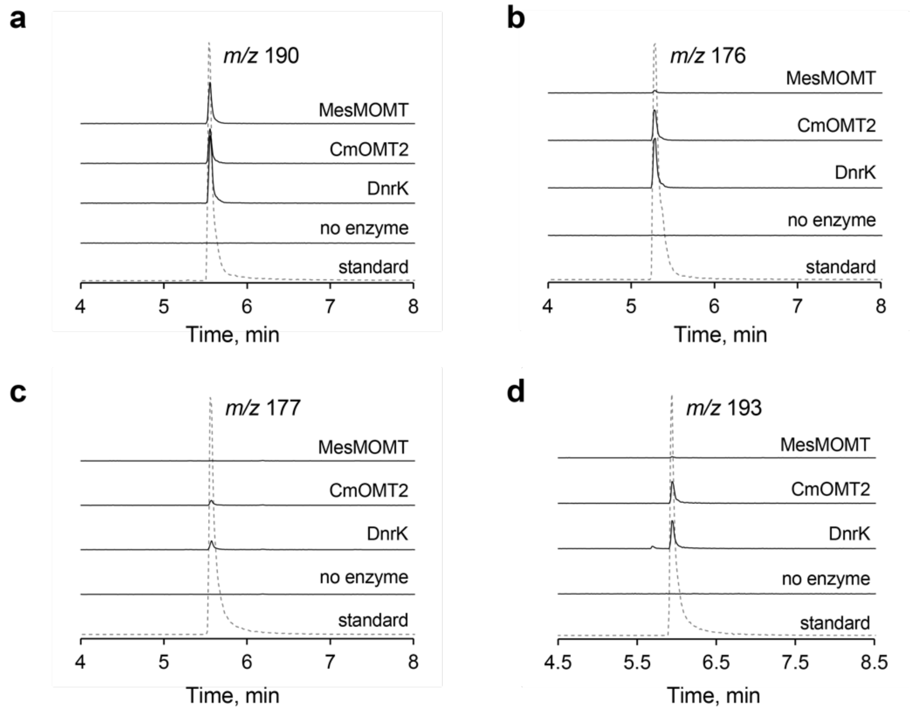

**Supplementary Figure 6.** Extracted ion chromatograms in low-resolution LC-MS of reaction products of DnrK, CmOMT2 and MesMOMT with substrates **12, 13, 16** and **18** compared to the "no enzyme" control and chemically synthesised reference compounds. a) Substrate **12**; b) substrate **13**; c) substrate **16**; d) substrate **18**.

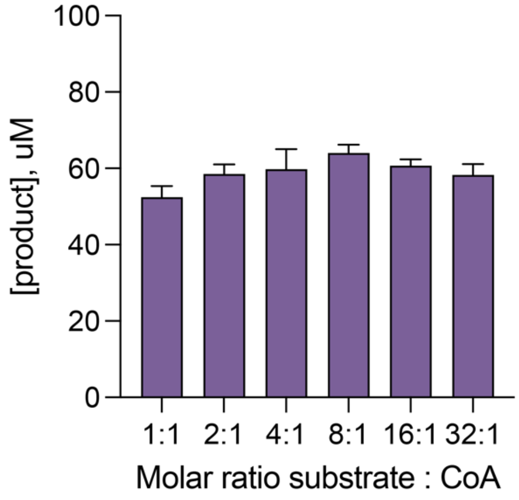

**Supplementary Figure 7.** Effect of decreasing CoA concentration on the efficiency of the cascade. The data are represented as mean ± SD; n=3.

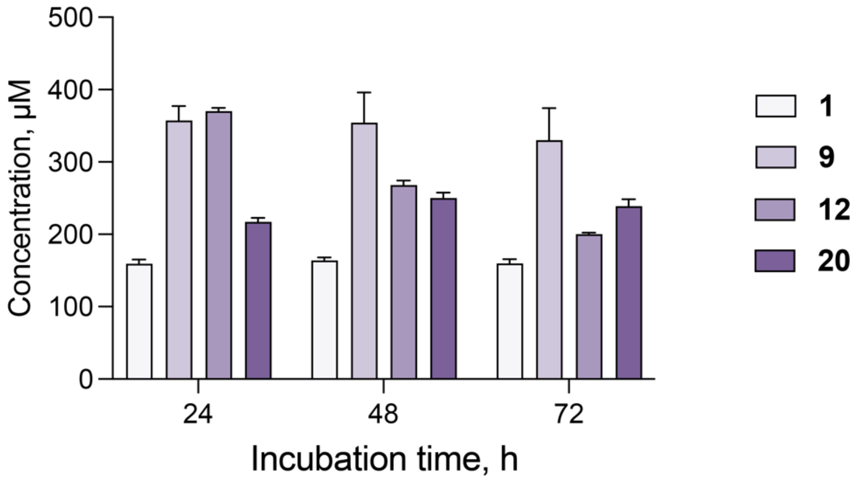

**Supplementary Figure 8.** Concentrations of cascade intermediates with increased substrate load (1000 µM). Apart from the initial concentration of **1**, all other conditions were kept constant. The data are represented as mean ± SD; n=3.

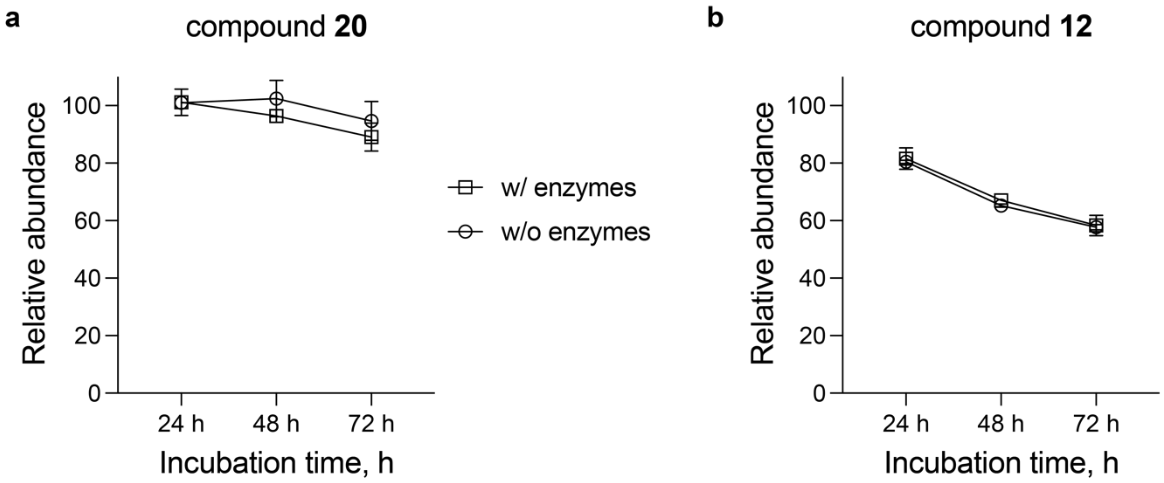

**Supplementary Figure 9.** Incubation of final product **20** (a) and quinolone intermediate **12** (b) in the reaction mix with or without the enzymes for up to 72 h. Relative Abundance was calculated by normalising the concentration of the compound at a particular timepoint relative to that at 0 h. The data are represented as mean ± SD; n=3.

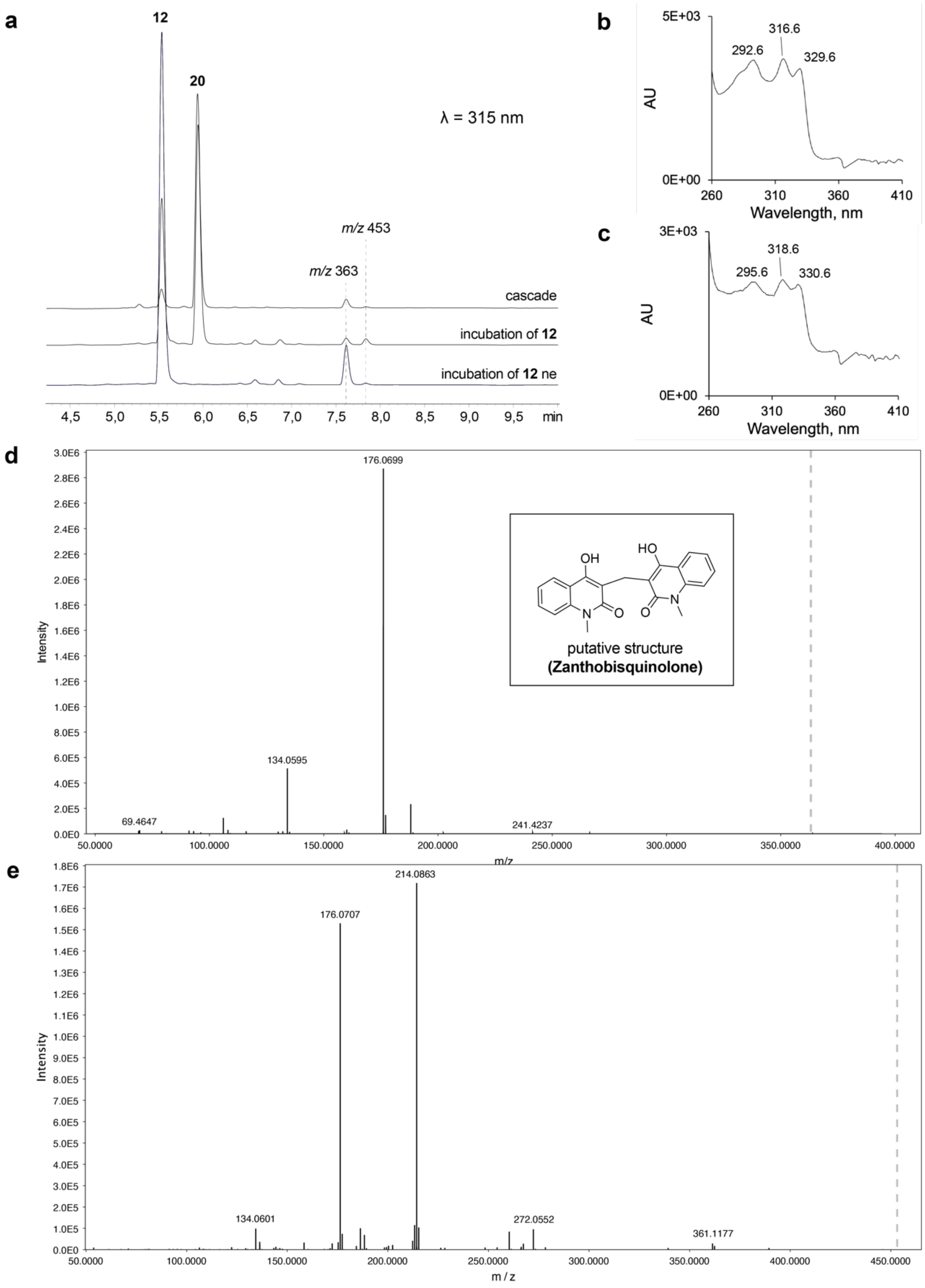

**Supplementary Figure 10.** Formation of side products from 12 and 20. a) UV chromatograms showing the formation of side products upon incubation of **12** in the reaction mix with or without the enzymes (ne) for 48 h. b) UV absorption spectrum of the side product with *m/z* 363. c) UV absorption spectrum of the side product with *m/z* 453. D) ESI-HR-MS/MS spectrum (positive mode) of the side product with *m/z* 363. Observed *m/z*: 363.1344; expected *m/z* of a dimer (zanthobisquinolone) = 363.1345, calculated for $[C_{21}H_{19}N_2O_4]^+$. e) ESI-HR-MS/MS spectrum (positive mode) of the side product with *m/z* 453. Observed *m/z*: 453.1484.

```
AthePKS         ------------------------MTTQPALCITGLGHQYPPYLMTPDKFDQ---LAARFH  34
AmQNS           MVTMEEIR-KAQRAEGLATILAISTATPPNCVI--QADYPDYYFKITNSEHMTELKEKFR  57
CmQNS           MESMAKVKNFLKRSKGPASILAIGTANPPTCFN--QSDYPDFYFRVTDCEHKTELKDKFK  58
RgACS           MESLKEMR-KAQKSEGPAAILAIGTATPDNVYI--QADYPDYYFKITKSEHMTELKDKFK  57
CmACS           MVTMEEIR-RAKRAEGLATILAISTATPPNCVI--QADYPDYYFGITNSEHMTELKEKFK  57
                                        *: *         :** : :   . ::    *   :*:

AthePKS         DTERPGLKKLIQINHTTGIESRASILPYTTGFATQSIPPTITEINHFFRSAGVDLAVSAC  94
AmQNS           RLCE---KSMIRKRHMCLTEEILKANPNMCLH----MGTSLNARQDISLVEVPKLGKEAA  110
CmQNS           RICD---RSAVKKRYLHVTEEVLKENPSMRS------YPSLDARQALLIEQVPKLGKEAA  109
RgACS           TLCE---KSMIRKRHMCFSQEFLKANPEVCKH----MGKSLNARQDIAVVETPRIGKEAA  110
CmACS           LLCE---KSMIRKRHMCLTEEILKANPNMCLY----MGTSLDARQDISLVEVPKLGKEAA  110
                       :. :: .:    :.  .  *            ::   : :       :. .*.

AthePKS         KKALRDARTPPAKITHTVAVTATNQGCPGYDLLVAKKLGLRLSSSSARPALDRTLLSGVG  154
AmQNS           TKAIKEWGQPKSKITHLIFCTSAGVDMPGADYQLTRLLGLS-------PEVKRMMIYQQG  163
CmQNS           AKAIKEWGQPLSKFTHLVFSAMAGVDIPGADLRLMNLLGLE-------PSVKRLMIYSQG  162
RgACS           VKAIKEWGHPKSSITHLIFCTSAGVDMPGADYQLTRMLGLN-------PSVKRMMIYQQG  163
CmACS           TKAIKEWGQPKSNITHLIFCTSAGVDMPGADYQLTRLIGLN-------PDVKRMMIYQQG  163
                 **:::   * :.:** :  : :. . ** *  : . :**        * :.* ::   *

AthePKS         CAGGLSVLRTAAQLACGASARGKPARILAYACELCMPNGRHDLAEVAREGDPDG--ISIV  212
AmQNS           CYAGATVLRLAKDLTEN----NKGSRVLIVCSENTVPTFR--------GPSDTHIDSLVG  211
CmQNS           CFIGGAAIRCAKDVAEN------GARVLVVFSDIVNMYFH--------EPQEAHLDILVG  208
RgACS           CYAGGTVLRLAKDLAEN----NKGSRVLVVCSELTAPTFR--------GPSPDAVDSLVG  211
CmACS           CYAGATILRLAKDLAEN----NKGSRVLVVCSENTIPTFR--------GPSYTHIDSLVG  211
                *  * : :* * ::: .       :*:*   .:      :          .       :

AthePKS         AALFSDAAGAVVLCNEVALAEDREGGADPDSEDEDEEAVRPVYRLLQWGHDVIPDTIEHM  272
AmQNS           QALFADGAAALIV------------GADP-----DASIERPLYHIVSASQTLLPDSDGAI  254
CmQNS           QAVFGDGAAAVIV------------GADP-----EVSIERPLFHVVSSTQMSVPDTNKFI  251
RgACS           QALFADGAAALVV------------GADP-----DTSVERALYYIVSASQMLLPDSDGAI  254
CmACS           QALFADGAAALIV------------GADP-----DTSIERPLYHIVSASQTLLPDSDGAI  254
                 *:*.*.*.*:::            ****     : .  * :: ::.  :  :**:   :

AthePKS         GFTTEGDGYHTTLTRAVPEHVKKAIGPIFSTLLPSYQQQLQSCTPETINKADQQLTIHDF  332
AmQNS           EGHIREAGLTVHLKKDVPAFFSANIEKSLVDAF----------TPIGIS------DWNSI  298
CmQNS           RAHVKEMGMELYLSKDVPATVGKNIEKLLVDAV----------SPFGIS------DWNSL  295
RgACS           EGHIREEGLTVHLKKDVPALFSANIDTPLVEAF----------RPLGIS------DWNSI  298
CmACS           EGHIREAGLTVHLKKDVPEFFSANIEKSLVDAF----------TPIGIS------DWNSI  298
                    .  *    *.: **  .   *   :   .           *  *.        :.:

AthePKS         EFALHPGGEAIIDGAKEVLGLTEDQIKVTRETYRTKGNSSSPTVLILLDKLRSEANRKR-  391
AmQNS           FWIAHPGGPAILDQVEEKLGLRKDKLKASRHVMSEFGNMSSACVLFILDEMRKTCLEEGK  358
CmQNS           FYSVHPGGRAILDQVELNLGLGKEKLRASRHVLSEYGNMGGSSVYFILDEIRKKSMQEAK  355
RgACS           FWIAHPGGPAILDQIEVKLGLKEDKLRASKHVMSEYGNMSSSCVLFVLDEMRNKSLQDGK  358
CmACS           FWIAHPGGPAILDQVEAKLGLKKDKLRASRHVMSEYGNMSSACVLFILDEMRNKCLEEGK  358
                 :  **** **:*  :  *** :::::.::..    ** ..  * ::**::*. . ..

AthePKS         -------ENVVAASFGPGMTIEMAMLRRCEDDE--    417
AmQNS           ATTGEGLDWGVLFGFGPGLTVETVVLRSVPIEA--    391
CmQNS           PTTGDGLEWGVLFAIGPGLTVETVILLSVPIDSAC    390
RgACS           STTGEGLDWGVLFGFGPGLTVETVVLRSVPVEA--    391
CmACS           ATTGEGLDWGVLFGFGPGLTVETVVLHSLPIEA--    391
                       :  *  .:***:*:* .:*     :
```

**Supplementary Figure 11.** Multiple sequence alignment of AthePKS with known quinolone and acridone synthases. Catalytic residues are highlighted in grey; residues of the substrate-binding pocket are highlighted in red; other residues of the active site cavity are highlighted in blue. AmQNS – quinolone synthase from *Aegle marmelos*[4]; CmQNS – quinolone synthase from *Citrus microcarpa*[5]; RgACS – acridone synthase from *Ruta graveolens*[6]; CmACS – acridone synthase from *Citrus microcarpa*[5].

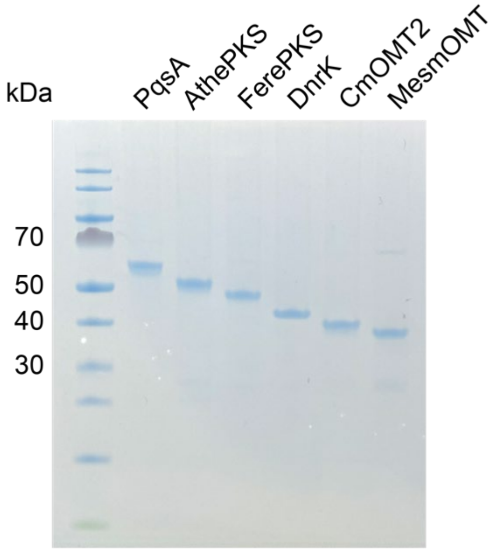

**Supplementary Figure 12.** SDS-PAGE of the purified proteins used for in vitro reactions in this study. Expected molecular weights: 57.9 kDa for PqsA, 46.5 kDa for AthePKS, 45.7 kDa for FerePKS, 40.3 kDa for DnrK, 40.5 kDa for CmOMT2, 37.5 kDa for MesMOMT.

#### [1]H NMR data of the synthesised reference compounds

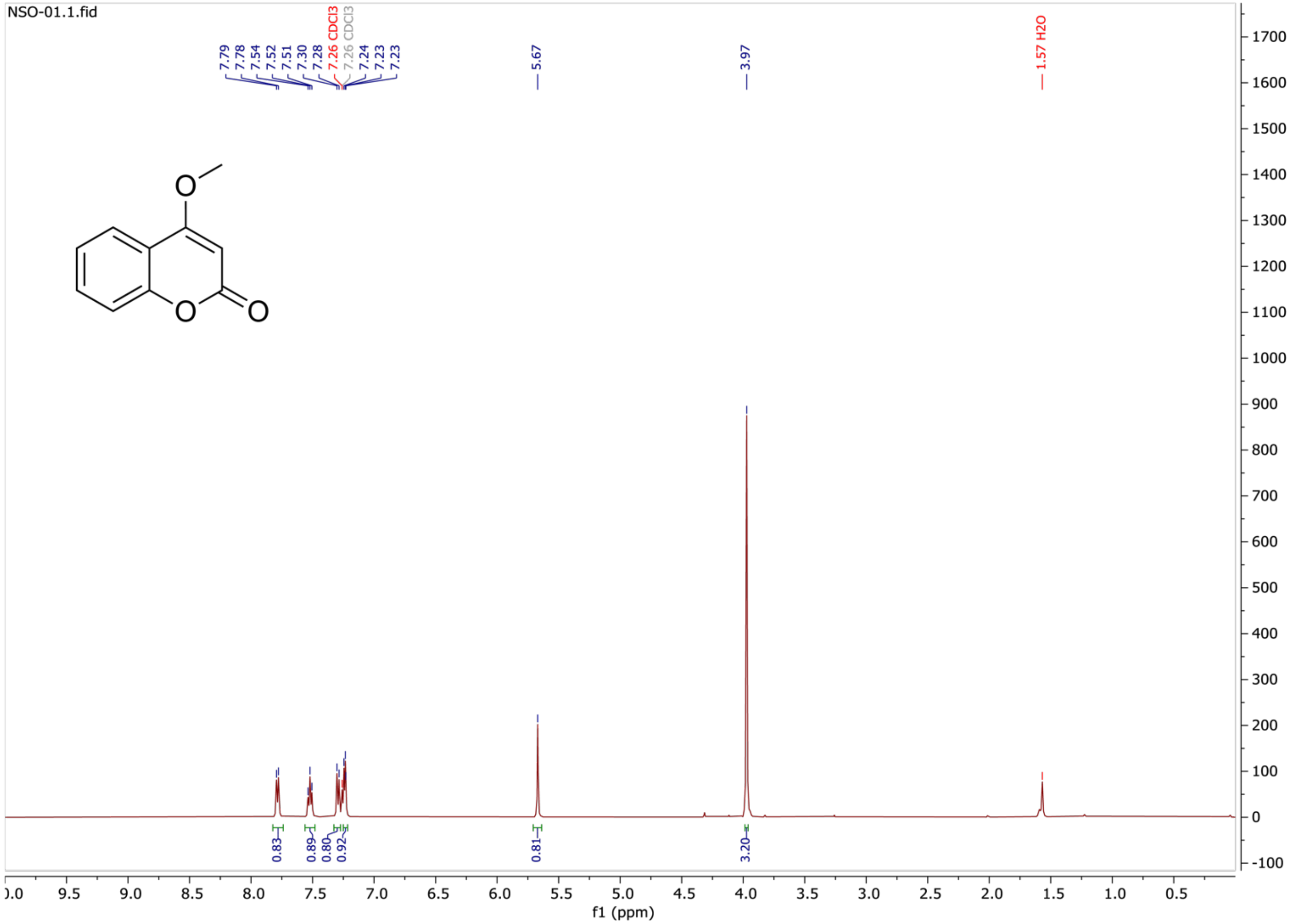

**Supplementary Figure 13.** $^1$H NMR spectrum of compound NSO-01.

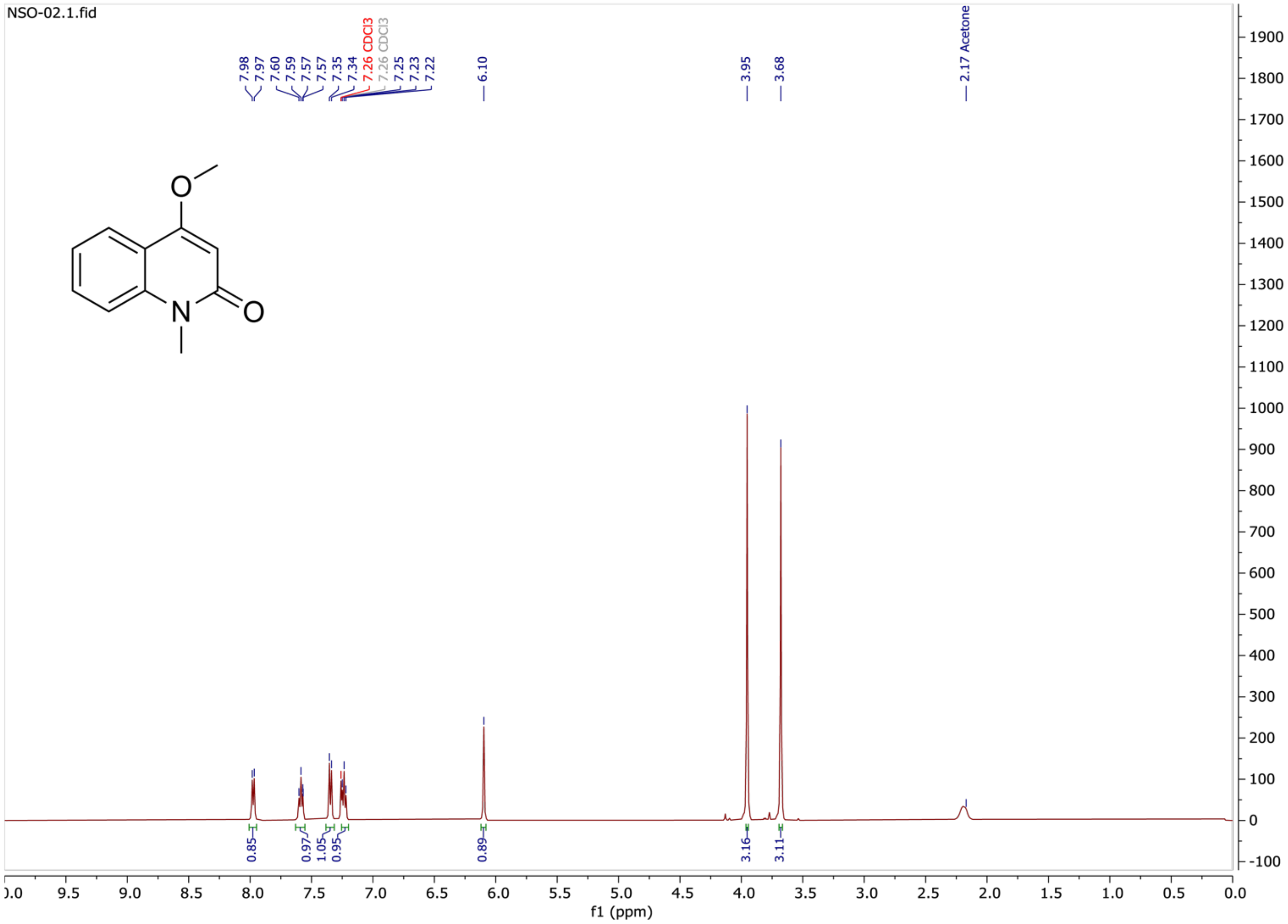

**Supplementary Figure 14.** $^{1}$H NMR spectrum of compound NSO-02.

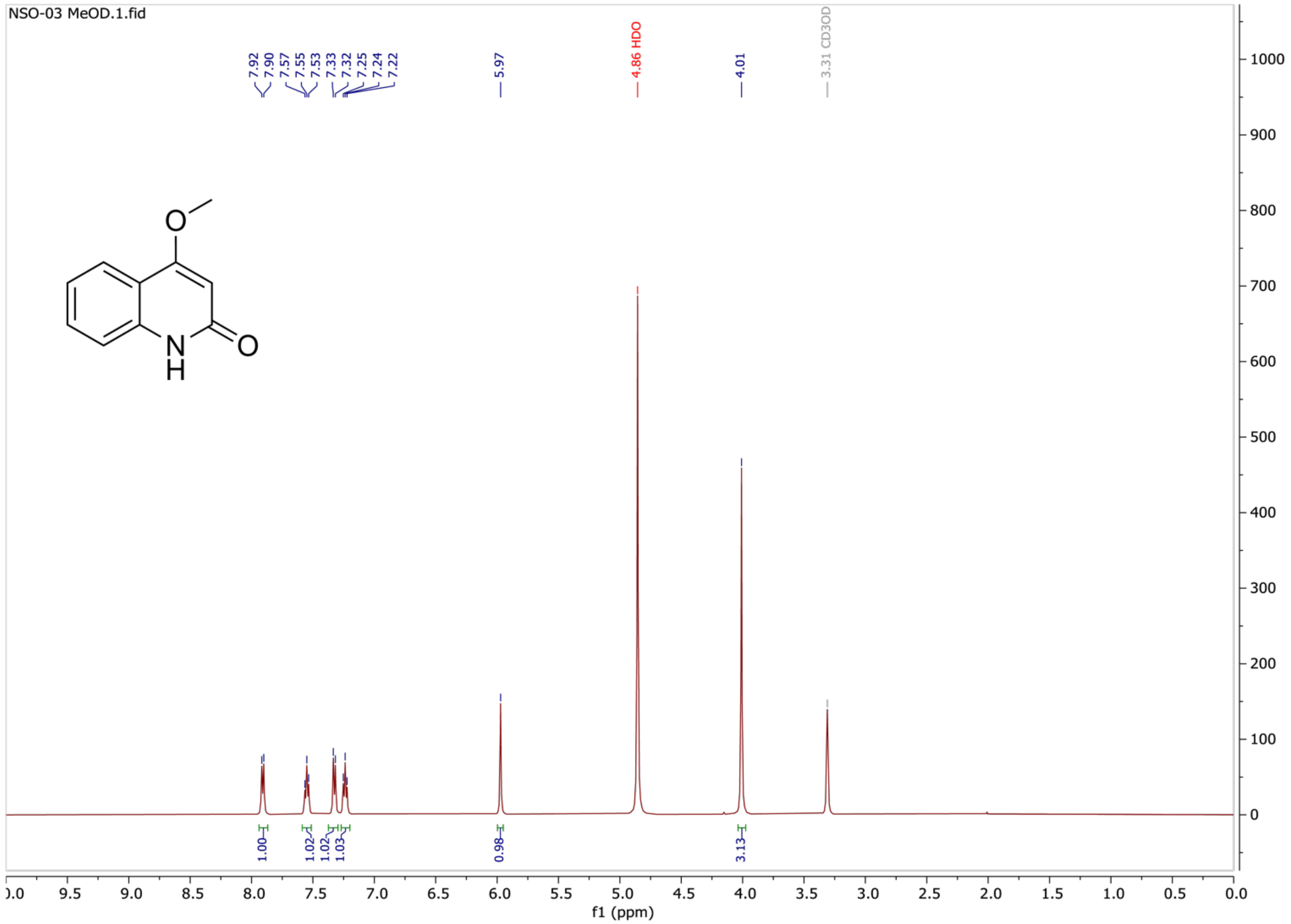

**Supplementary Figure 15.** $^{1}$H NMR spectrum of compound NSO-03.

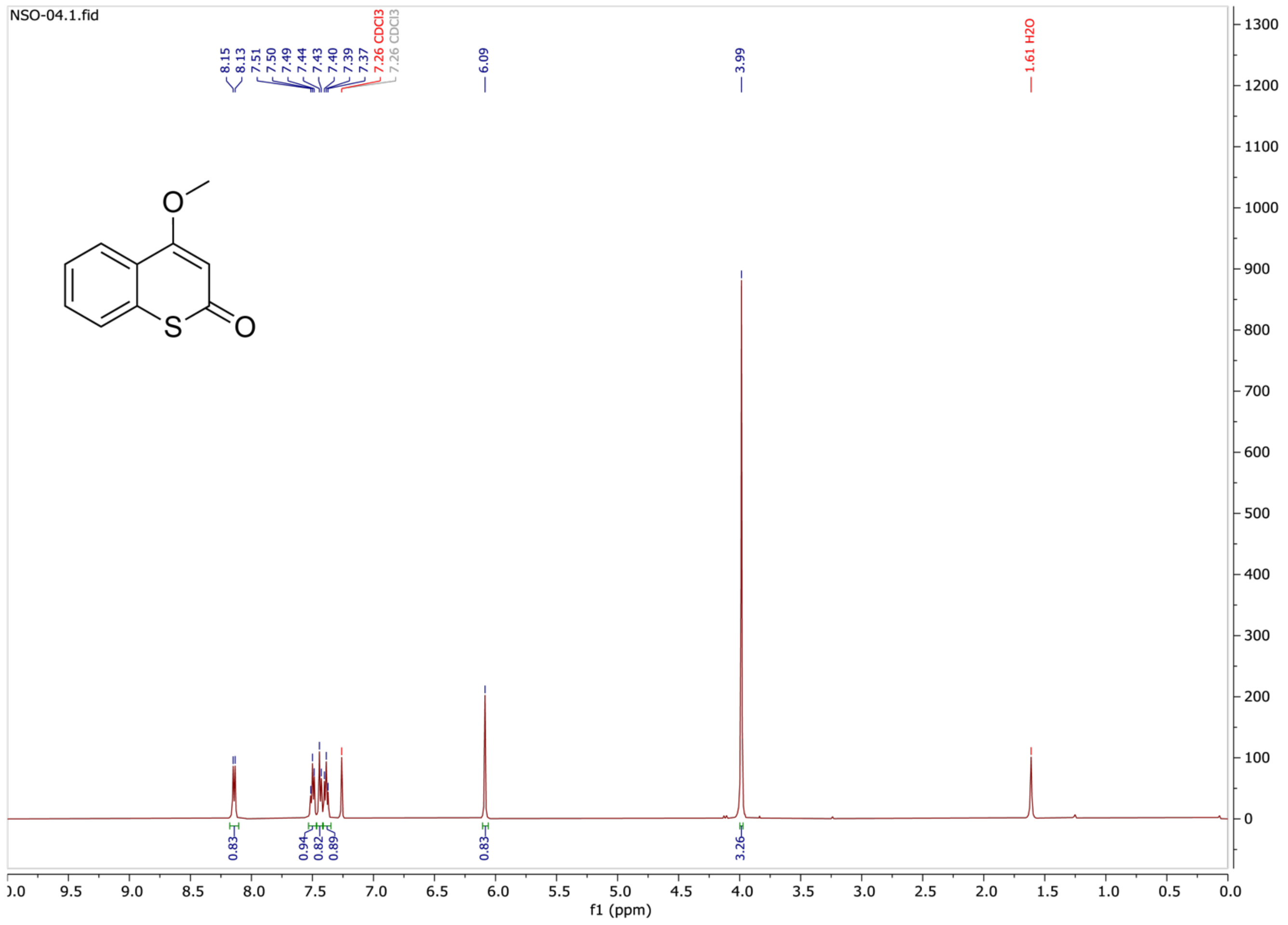

**Supplementary Figure 16.** $^{1}$H NMR spectrum of compound NSO-04.

#### Spectral data of all enzymatic products and reference compounds

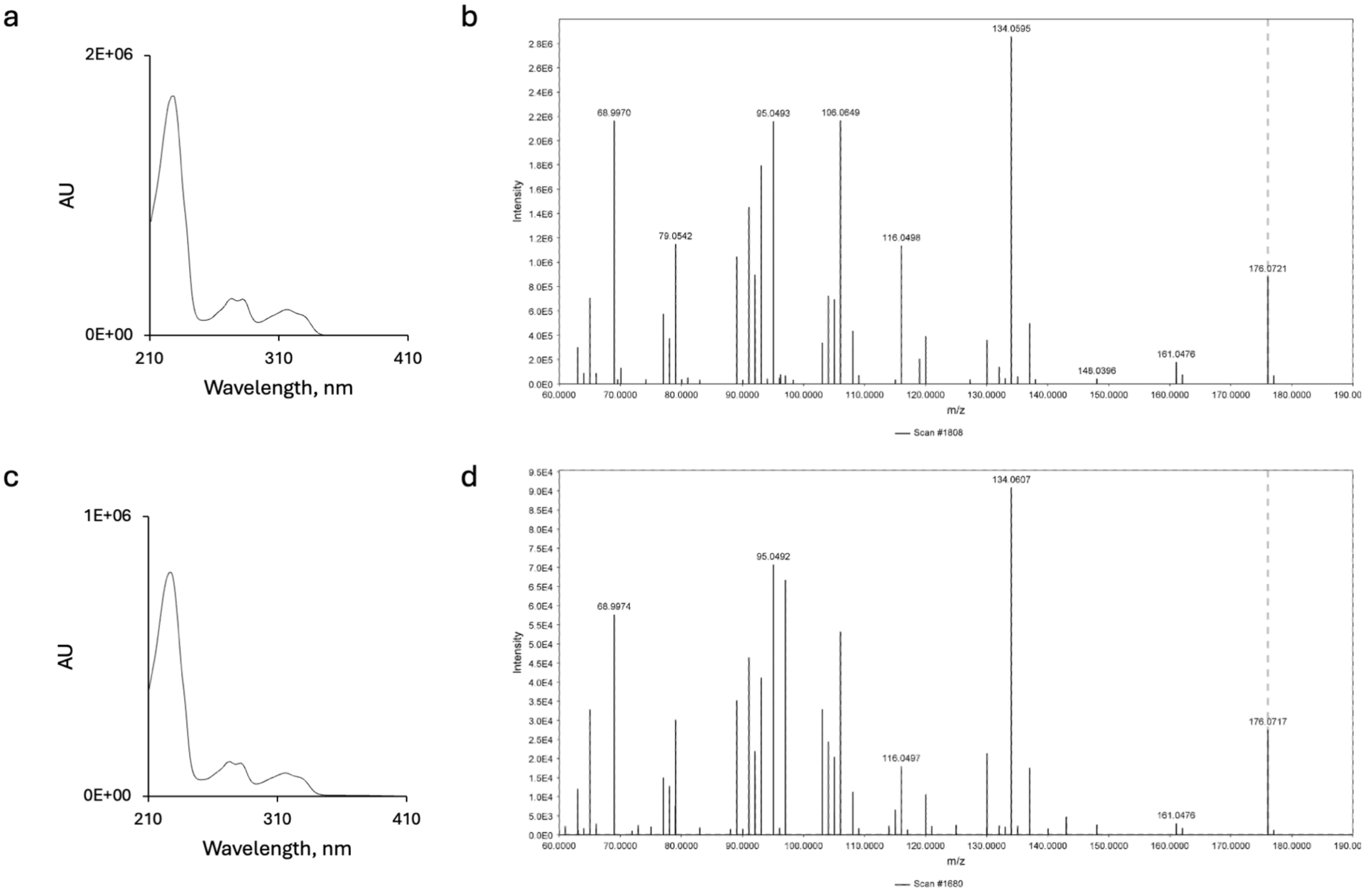

**Supplementary Figure 17.** Comparison of the UV absorption and ESI-HR-MS/MS spectra (positive ion mode) for a-b) reference compound **12** and c-d) enzymatically generated **12**. Observed *m/z* range: 176.0721 to 176.0717 (theoretical *m/z* = 176.0712, calculated for $[C_{10}H_{10}NO_2]^+$). The precursor ion is indicated with a dashed grey line.

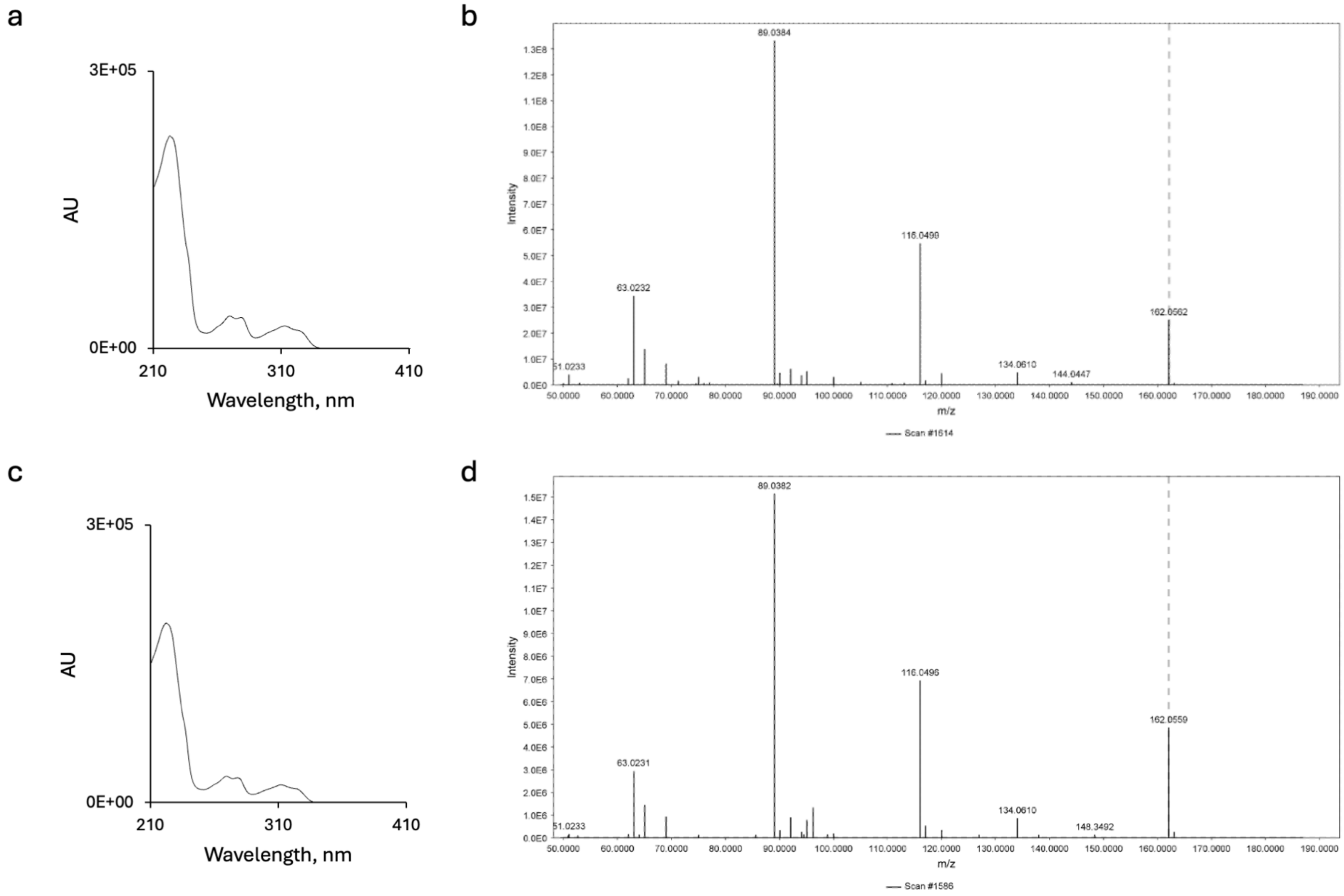

**Supplementary Figure 18.** Comparison of the UV absorption and ESI-HR-MS/MS spectra (positive mode) for a-b) reference compound **13** and c-d) enzymatically generated **13**. Observed *m/z* range: 162.0559 to 162.0562 (theoretical *m/z* = 162.0555, calculated for $[C_9H_8NO_2]^+$). The precursor ion is indicated with a dashed grey line.

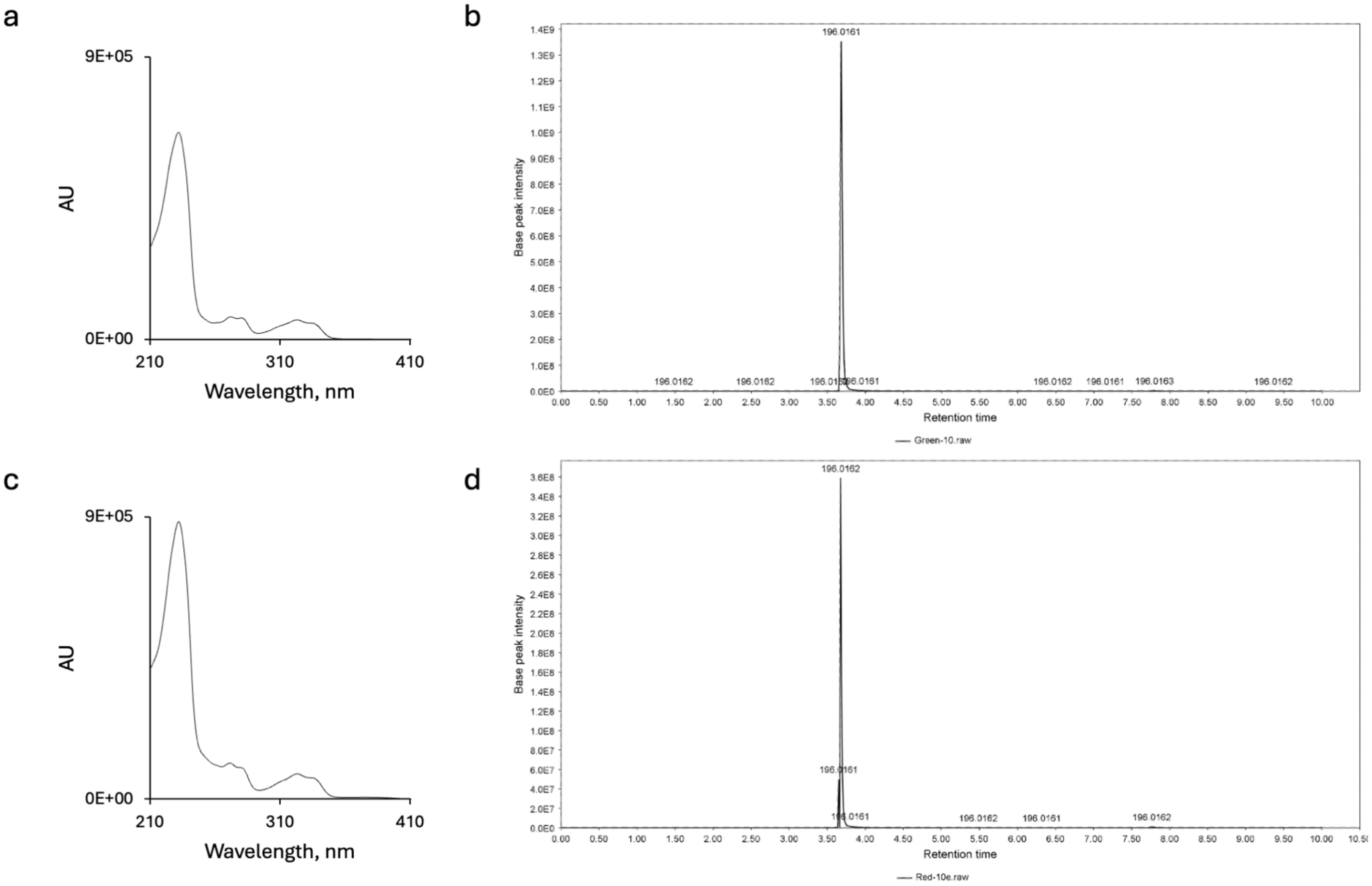

**Supplementary Figure 19.** Comparison of the UV absorption and extracted ESI-HR-MS spectra (positive mode) for a-b) reference compound **14** and c-d) enzymatically generated **14**. MS/MS fragmentation was not observed for compound **14**. Observed *m/z* range: 196.0161 to 196.0162 (theoretical *m/z* = 196.0165, calculated for $[C_9H_7ClNO_2]^+$).

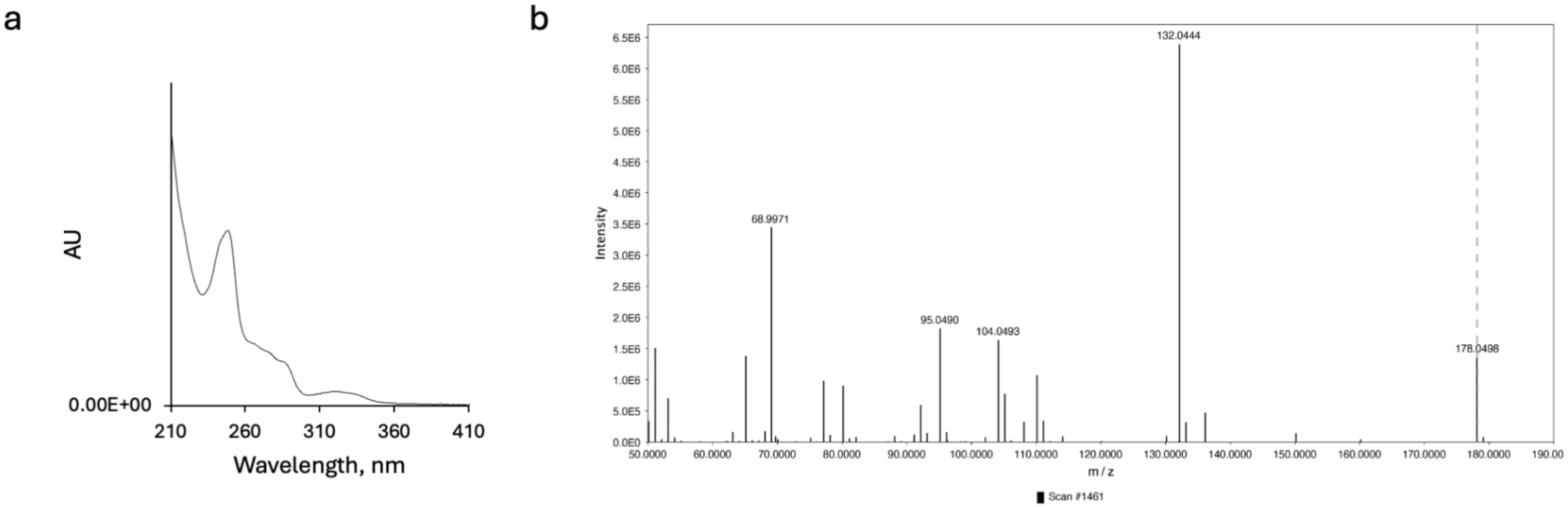

**Supplementary Figure 20.** UV absorption and ESI-HR-MS/MS spectra (positive mode) for a-b) enzymatically generated **15**. Observed *m/z* range: 178.0498 to 178.0500 (theoretical *m/z* = 178.0504, calculated for $[C_9H_8NO_3]^+$). The precursor ion is indicated with a dashed grey line.

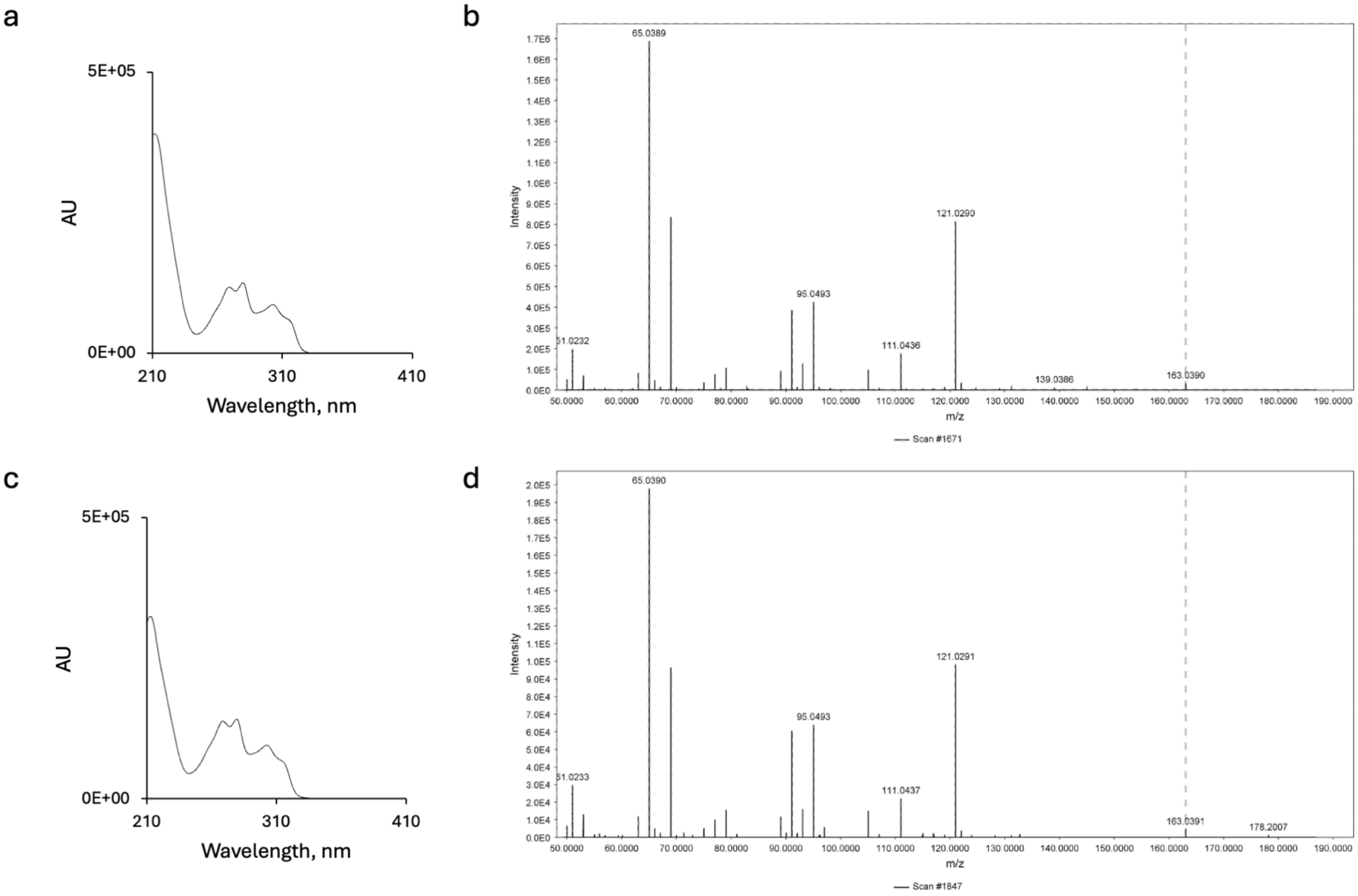

**Supplementary Figure 21.** Comparison of the UV absorption and ESI-HR-MS/MS spectra (positive mode) for a-b) reference compound **16** and c-d) enzymatically generated **16**. Observed *m/z* range: 163.0390 to 163.0391 (theoretical *m/z* = 163.0395, calculated for $[C_9H_7O_3]^+$). The precursor ion is indicated with a dashed grey line.

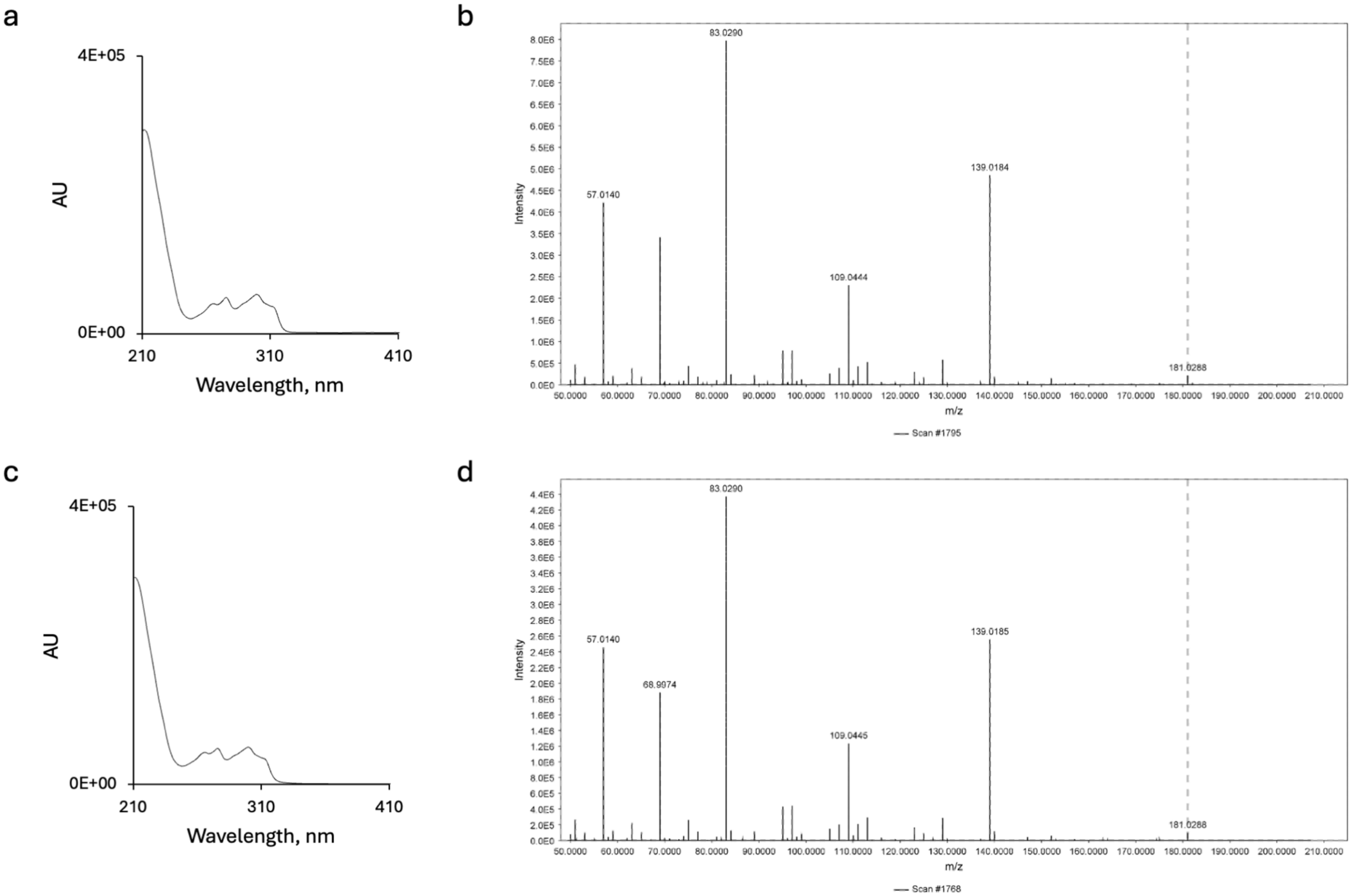

**Supplementary Figure 22.** Comparison of the UV absorption and ESI-HR-MS/MS spectra (positive mode) for a-b) reference compound **17** and c-d) enzymatically generated **17**. Observed *m/z*: 181.0288 (theoretical *m/z* = 181.0301, calculated for $[C_9H_6FO_3]^+$). The precursor ion is indicated with a dashed grey line.

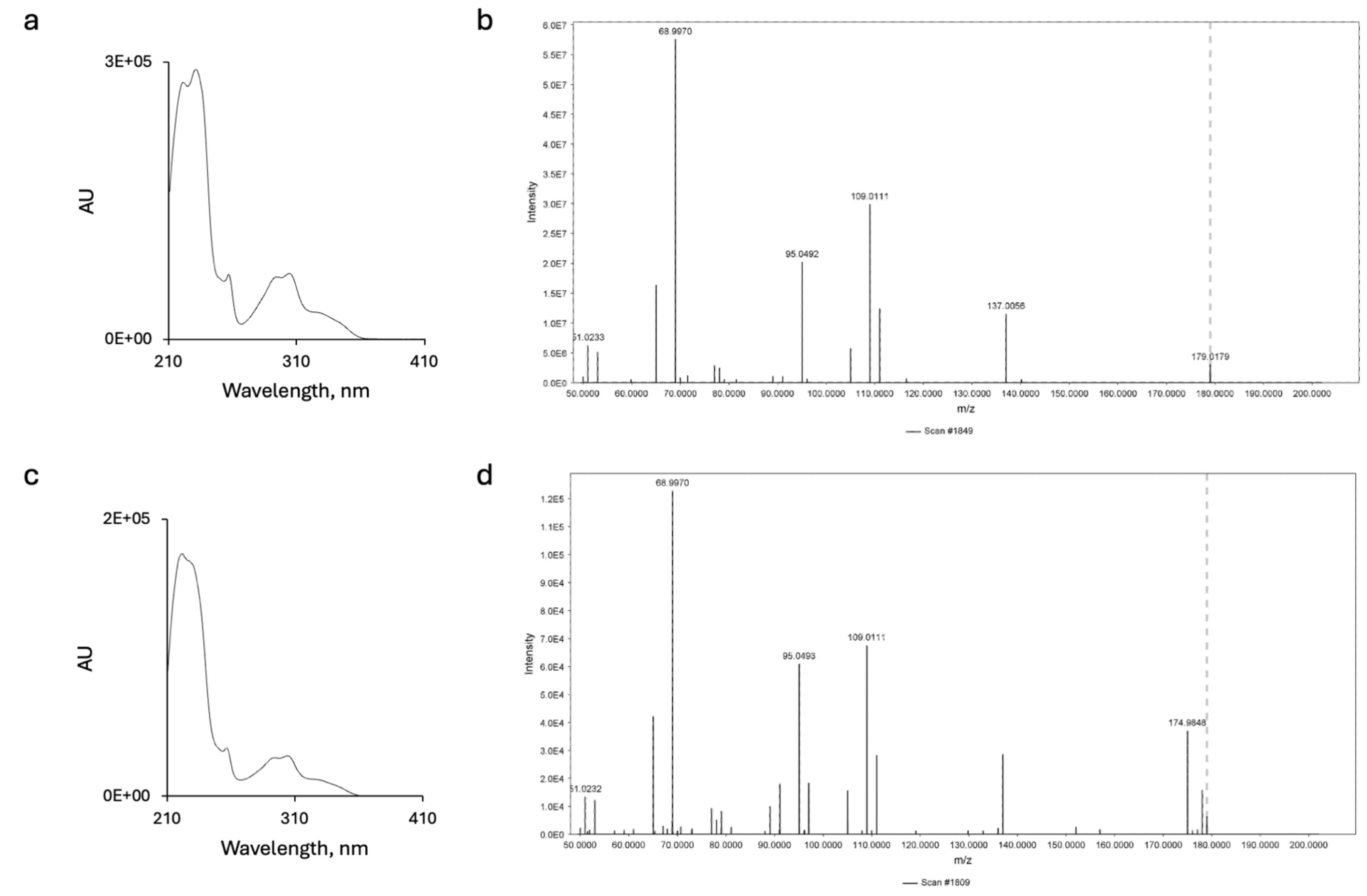

**Supplementary Figure 23.** Comparison of the UV absorption and ESI-HR-MS/MS spectra (positive mode) for a-b) reference compound **18** and c-d) enzymatically generated **18**. Observed *m/z* range: 179.0165 to 179.0179 (theoretical *m/z* = 179.0167, calculated for $[C_9H_7O_2S]^+$). The precursor ion is indicated with a dashed grey line.

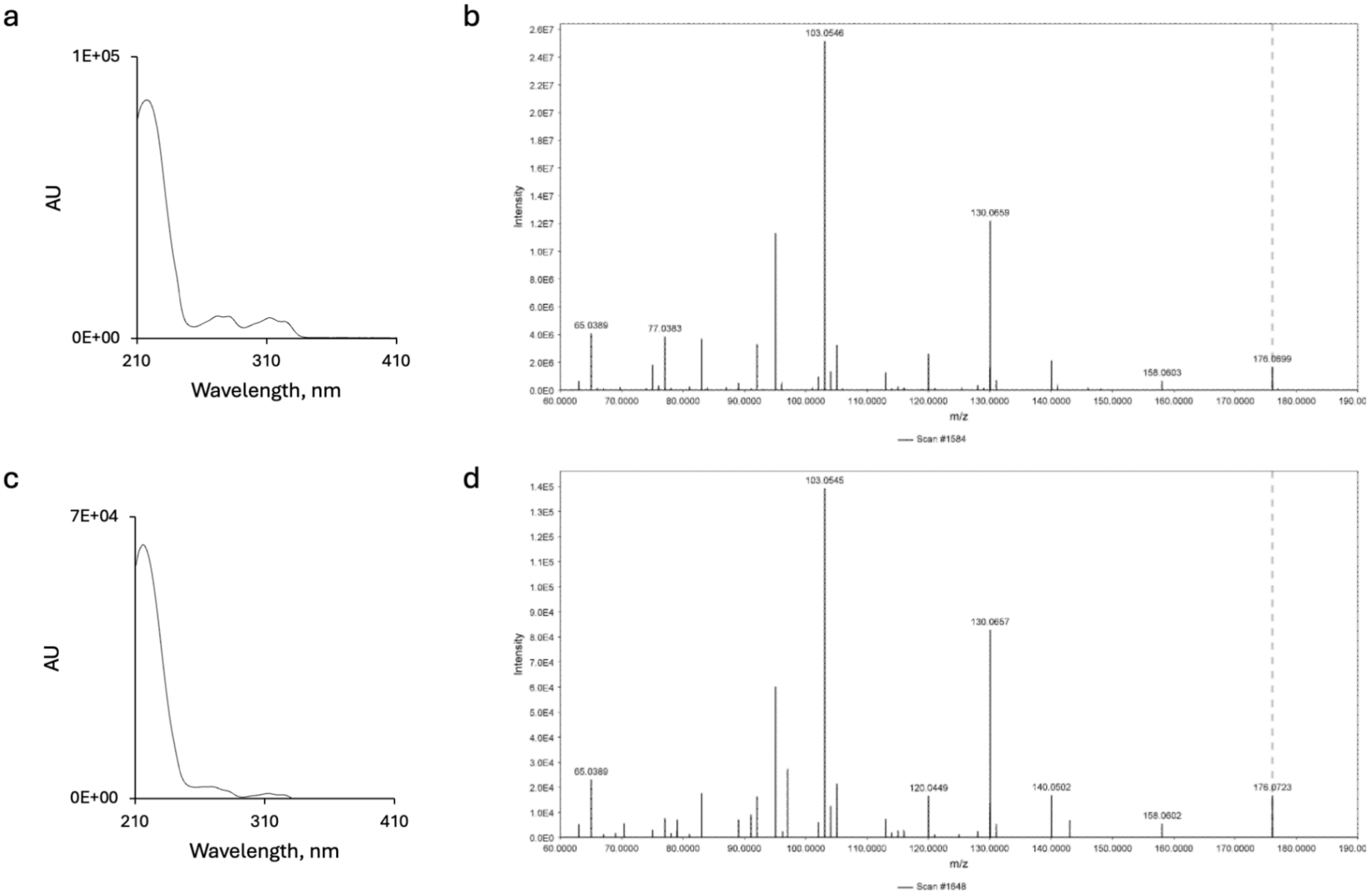

**Supplementary Figure 24.** Comparison of the UV absorption and ESI-HR-MS/MS spectra (positive mode) for a-b) reference compound **19** and c-d) enzymatically generated **19**. Observed *m/z* range: 176.0699 to 176.0723 (theoretical *m/z* = 176.0712, calculated for $[C_{10}H_{10}NO_2]^+$). The precursor ion is indicated with a dashed grey line.

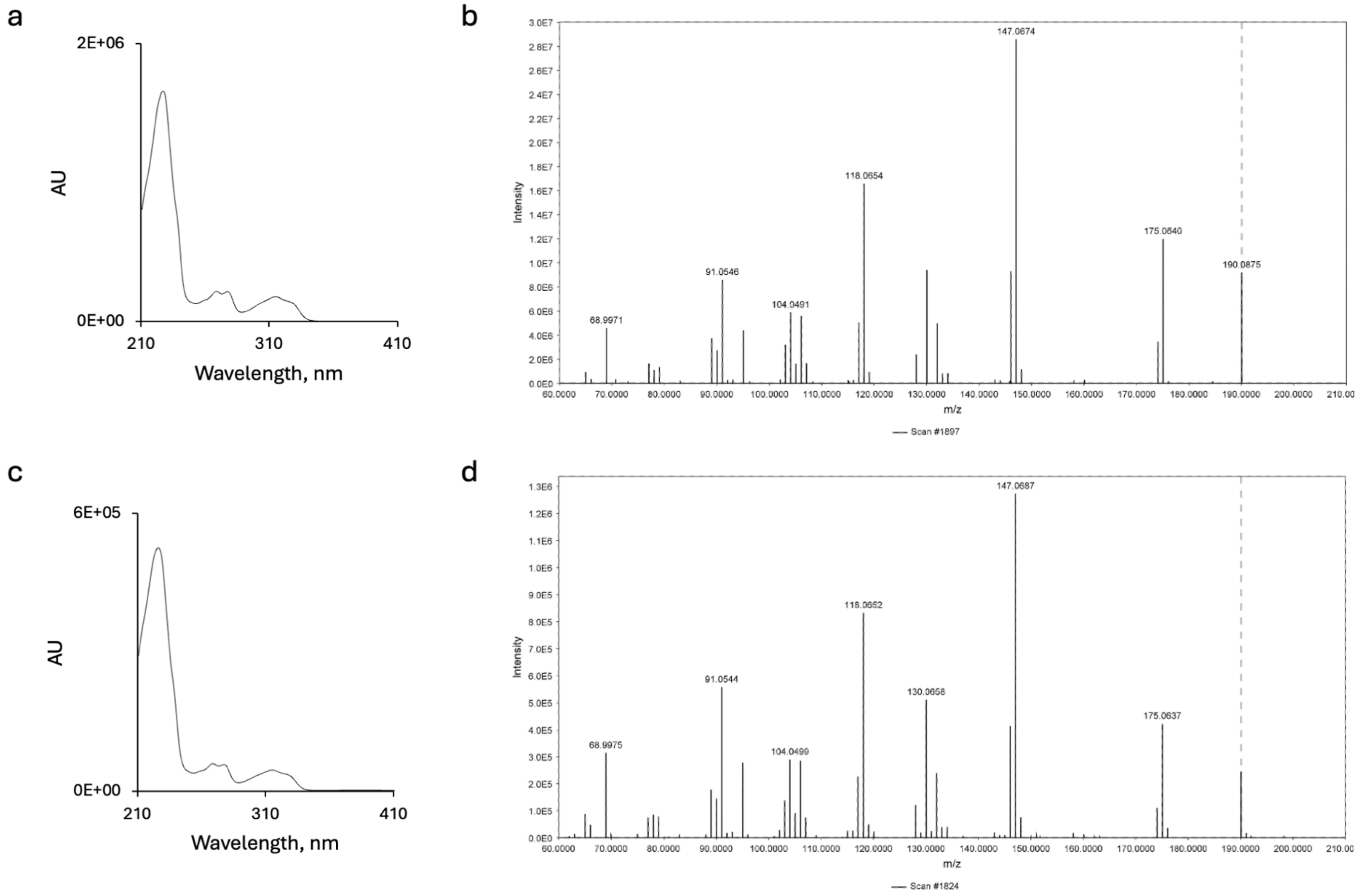

**Supplementary Figure 25.** Comparison of the UV absorption and ESI-HR-MS/MS spectra (positive mode) for a-b) chemically synthesised **20** (NSO-02) and c-d) enzymatically generated **20**. Observed *m/z* range: 190.0866 to 190.0875 (theoretical *m/z* = 190.0868, calculated for $[C_{11}H_{12}NO_2]^+$). The precursor ion is indicated with a dashed grey line.

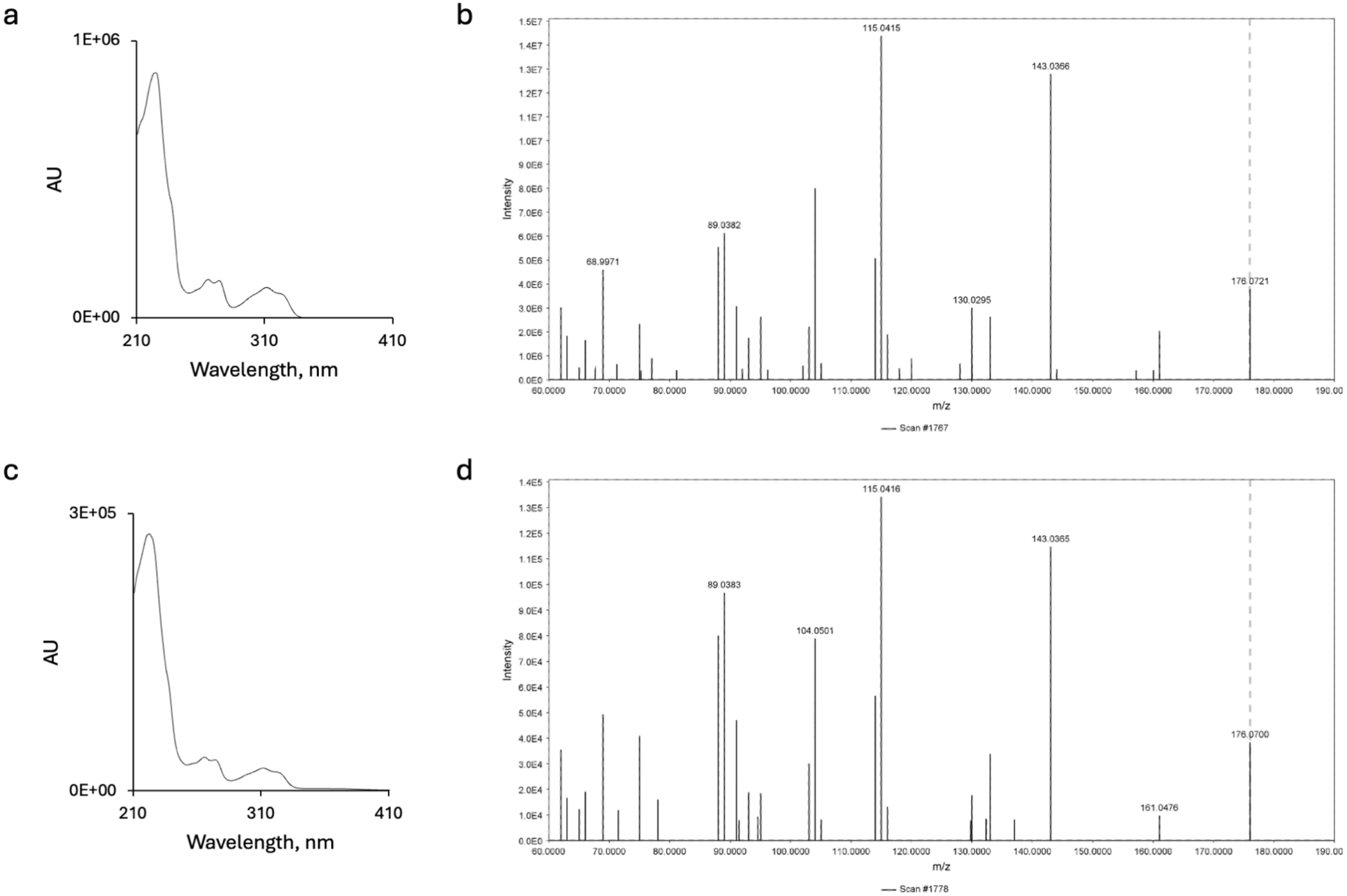

**Supplementary Figure 26.** Comparison of the UV absorption and ESI-HR-MS/MS spectra (positive mode) for a-b) chemically synthesised **21** (NSO-03) and c-d) enzymatically generated **21**. Observed *m/z* range: 176.0700 to 176.0721 (theoretical *m/z* = 176.0712, calculated for $[C_{10}H_{10}NO_2]^+$). The precursor ion is indicated with a dashed grey line.

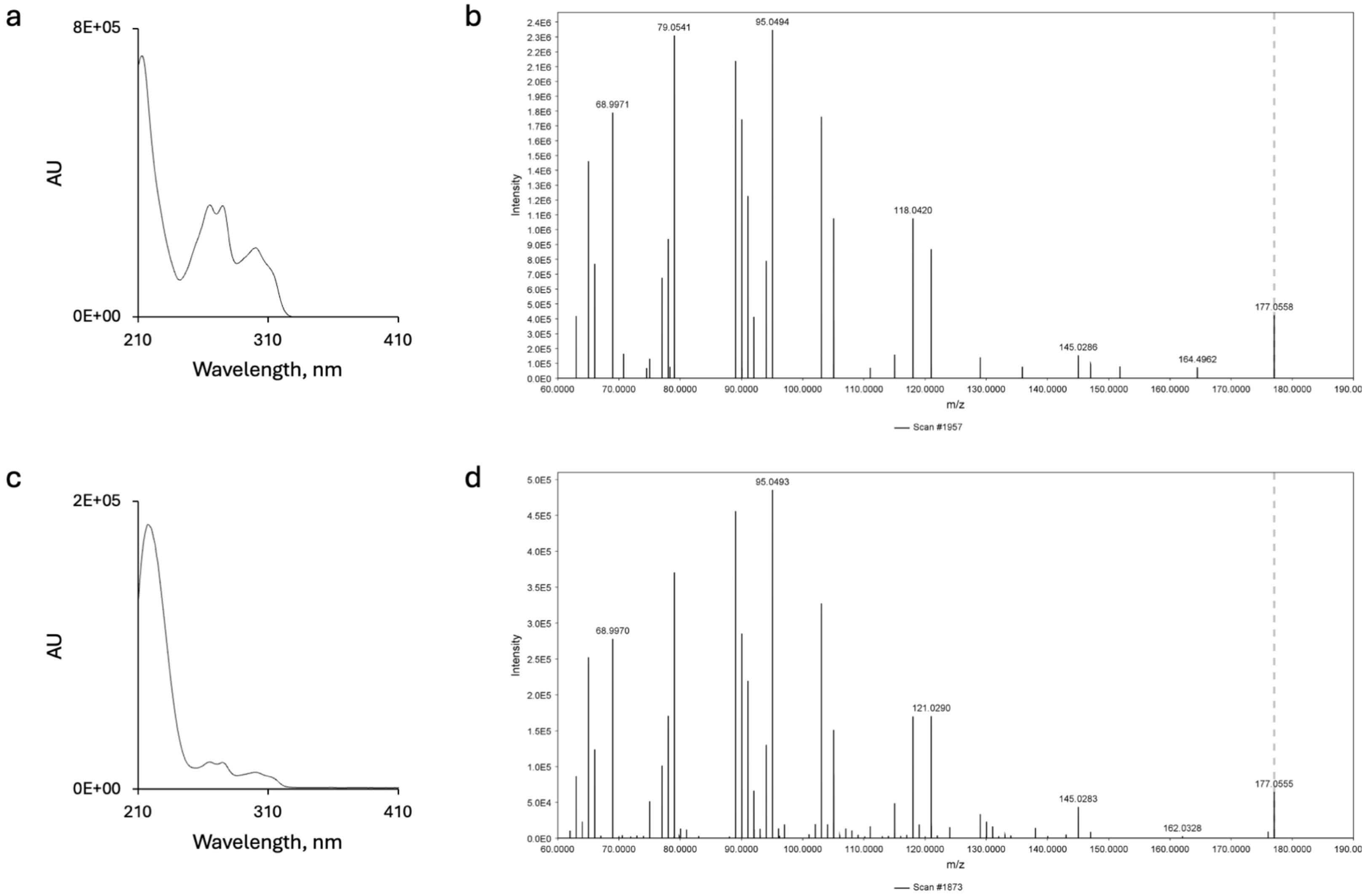

**Supplementary Figure 27.** Comparison of the UV absorption and ESI-HR-MS/MS spectra (positive mode) for a-b) chemically synthesised **22** (NSO-01) and c-d) enzymatically generated **22**. Observed *m/z* range: 177.0555 to 177.0558 (theoretical *m/z* = 177.0552, calculated for $[C_{10}H_9O_3]^+$). The precursor ion is indicated with a dashed grey line.

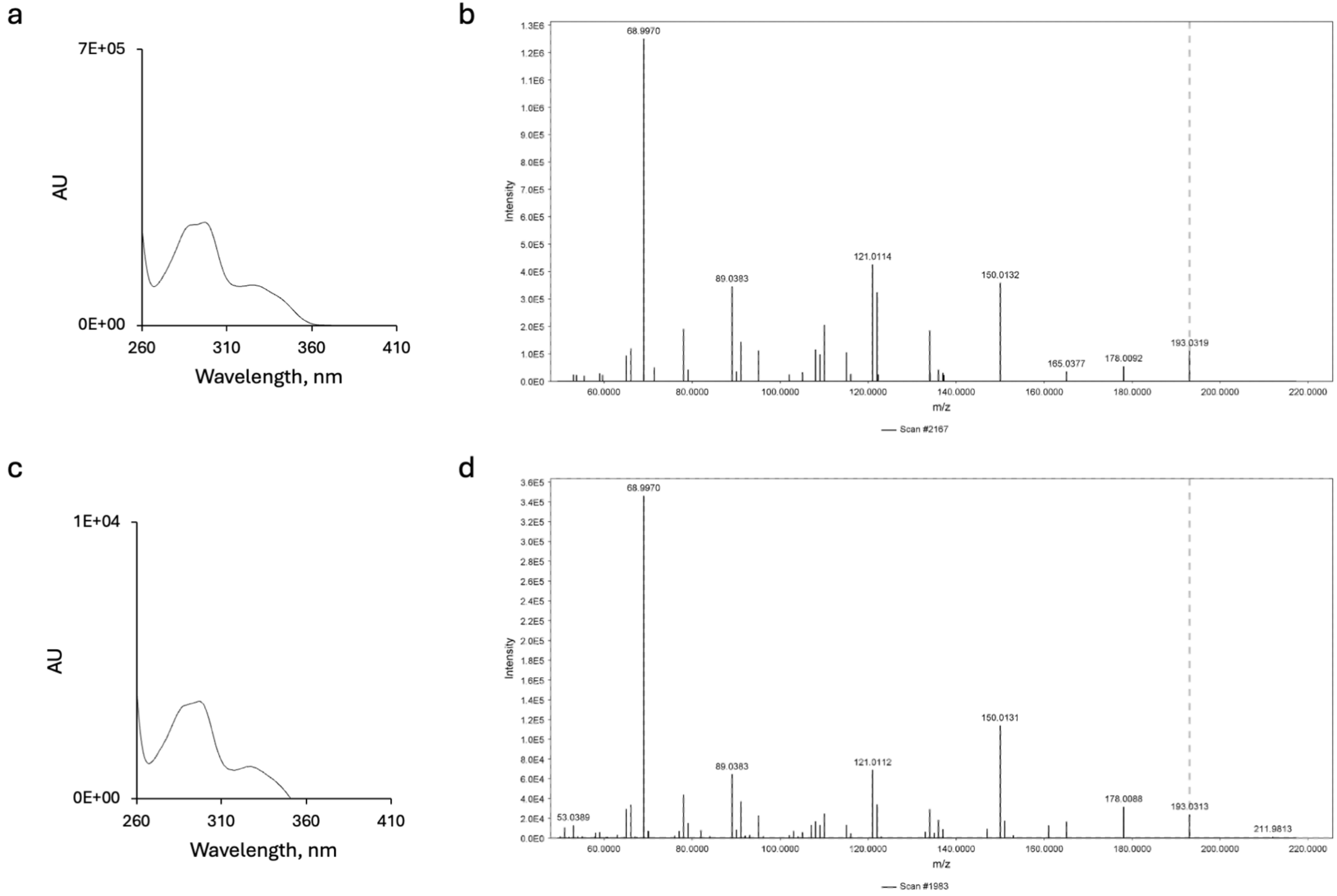

**Supplementary Figure 28.** Comparison of the UV absorption and ESI-HR-MS/MS spectra (positive mode) for a-b) chemically synthesised **23** (NSO-04) and c-d) enzymatically generated **23**. Observed *m/z* range: 193.0313 to 193.0319 (theoretical *m/z* = 193.0323, calculated for $[C_{10}H_9O_2S]^+$). The precursor ion is indicated with a dashed grey line.

#### Calibration plots

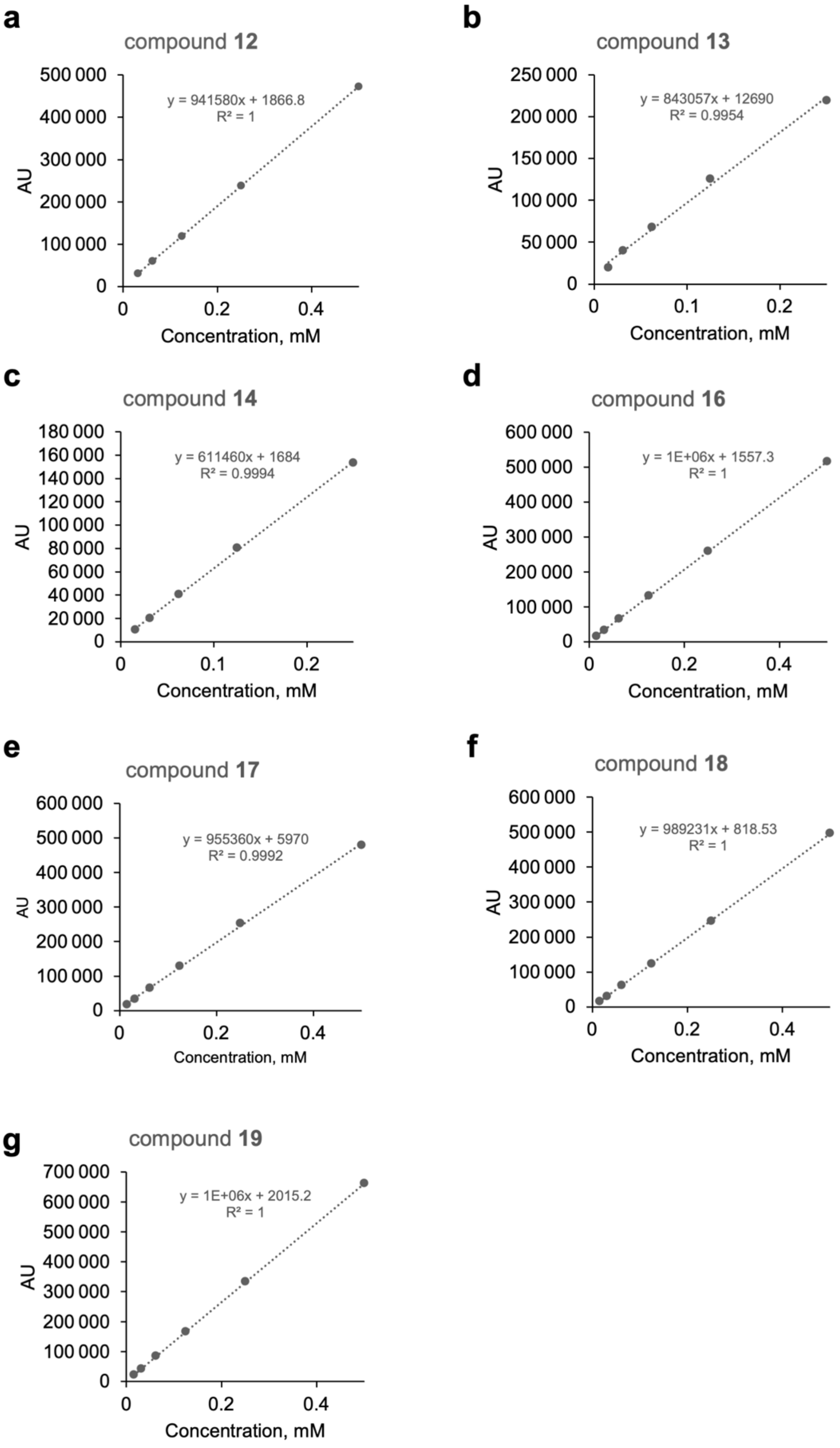

**Supplementary Figure 29.** Calibration plots used during Athe/FerePKS substrate scope analysis. a) Compound **12**; b) compound **13**; c) compound **14**; d) compound **16**; e) compound **17**; f) compound **18**; g) compound **19**. All compounds were dissolved in water : acidic methanol (1 : 1) and analysed by HPLC.

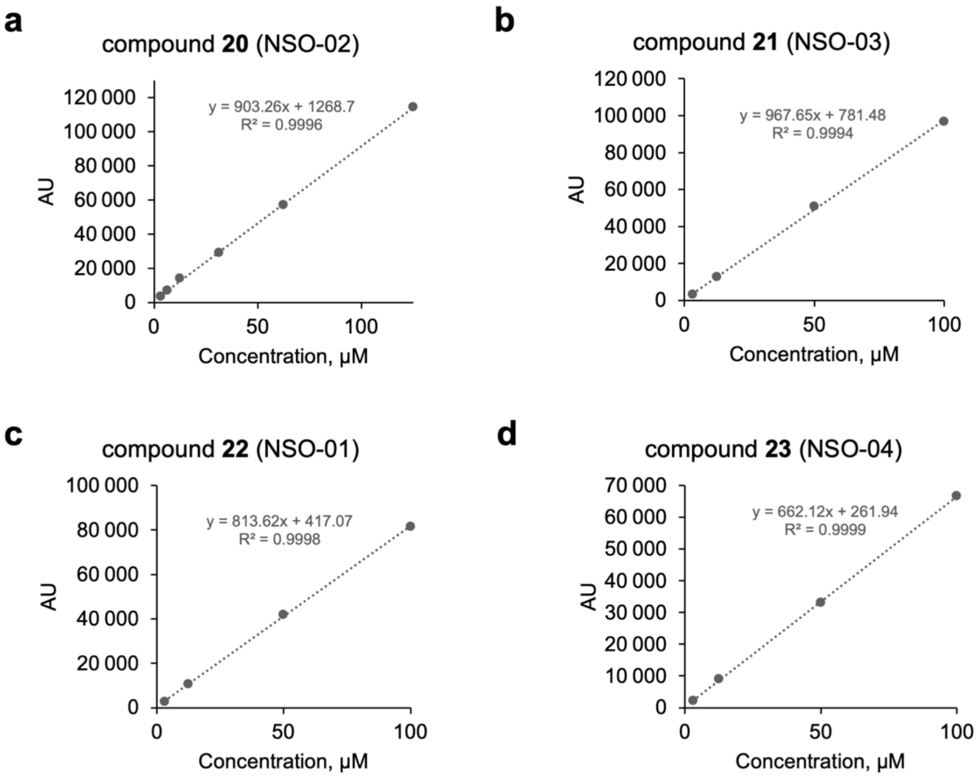

**Supplementary Figure 30.** Calibration plots used during OMT substrate scope analysis. a) Compound **20**; b) compound **21**; c) compound **22**; d) compound **23**. All compounds were dissolved in water : acidic methanol (1 : 1) and analysed by HPLC.

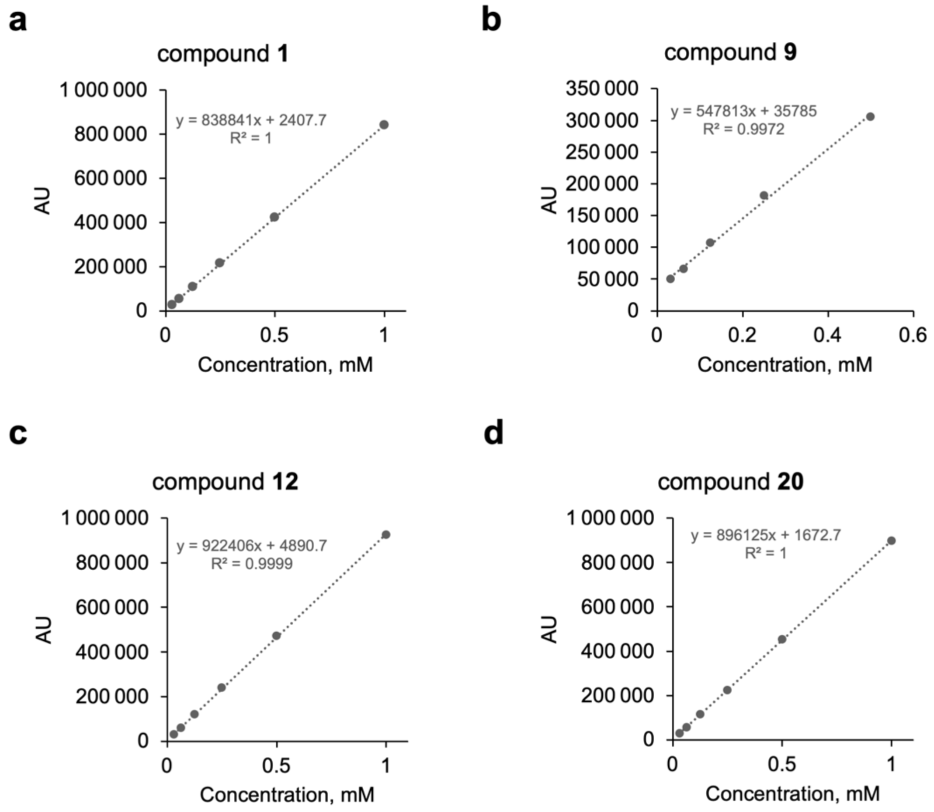

**Supplementary Figure 31.** Calibration plots used during cascade optimisation experiments. a) Compound **1**; b) compound **9**; c) compound **12**; d) compound **20**. All compounds were dissolved in water : acidic methanol (1 : 1) and analysed by HPLC.
